# Transient focal inactivation of the primary visual cortex abolishes saccadic inhibition

**DOI:** 10.64898/2026.03.06.710115

**Authors:** Tatiana Malevich, Yue Yu, Matthias P. Baumann, Xiyao Yu, Tong Zhang, Masatoshi Yoshida, Tadashi Isa, Ziad M. Hafed

**Author notes:** Corresponding authors: Ziad M. Hafed, Masatoshi Yoshida, Tadashi Isa. Equal contributions (Tatiana Malevich, Yue Yu, Matthias P. Baumann).

## Abstract

Most organisms are perpetually engaged in an active-perception cycle: at any moment, planning, executing, or recovering from a given own movement. In the oculomotor system, such a cycle is under a powerful, reflexive, and short-latency interruptive influence by exogenous sensory stimulation, but the multitude of possible pathways eventually converging on the final oculomotor control gateway make it unknown how reflexive oculomotor inhibition takes place. Here we show that transient focal inactivation of the primary visual cortex abolishes visually-driven saccadic inhibition, suggesting an unequivocal dominance of the geniculostriate visual pathway over this robust behavioral reflex. Using chronic lesions, computational modeling, and fine-grained eye-movement analyses, we then show that latent visual signals bypassing the geniculostriate pathway are still present, but may not be as central as previously anticipated. These results recast classic views on visual-motor reflexes, and motivate functional and theoretical investigations of the multiplicity of parallel input-output loops in the brain.

## Introduction

A hallmark of brain organization is massive parallelism and the presence of multiple hierarchies of nested anatomical loops. For example, in the primate visual system, besides the geniculostriate pathway from the retina to the cortex (1–4), direct retinal projections to structures like the superior colliculus (SC) (5–19) and pulvinar (20, 21) also exist (Fig. 1A).

**Figure 1.**
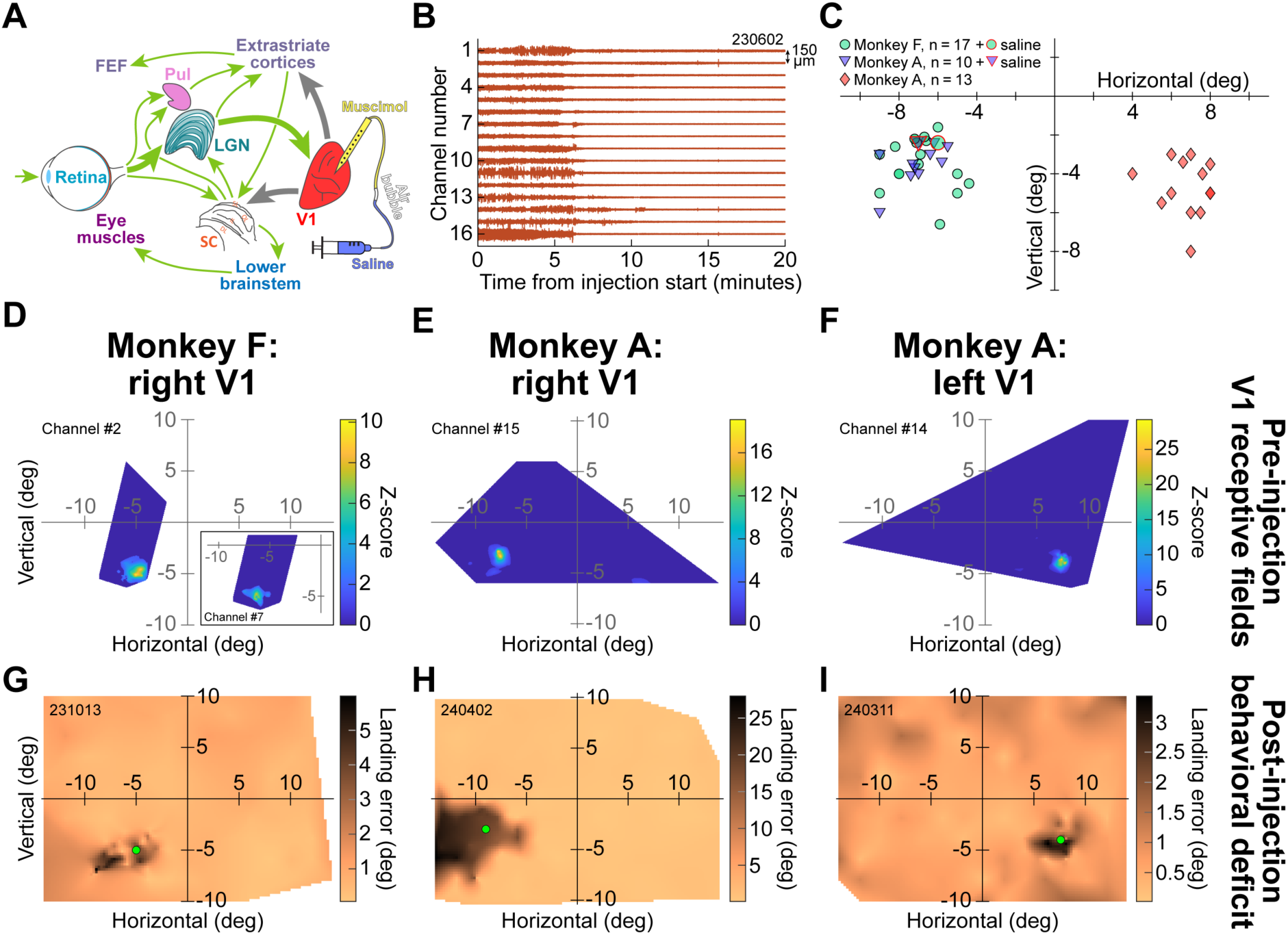
Reversible V1 inactivation caused localized visual scotomas. **(A)** We reversibly inactivated V1 via muscimol injection; this blocked V1 outputs to other brain areas, but visual signals could still reach the oculomotor system via alternative pathways (thin arrows show examples of such pathways). Pul: pulvinar; other acronyms are defined in the text. Brain areas not drawn to scale. **(B)** Example cortical activity as a function of time from muscimol injection start in one session (monkey F). The figure shows multi-unit activity in different channels along the depth of the injectrode. After a few minutes of injection, cortical spiking was largely lost. **(C)** Visual field locations targeted by our injectrodes across experiments. In monkey A, we tested both right and left V1. In each monkey, we also had a control saline injection at a location similar to the muscimol injection locations. **(D)** Example multi-unit visual receptive field (RF) from one sample session in monkey F. These data were obtained by presenting visual stimuli at different locations (Methods) but prior to muscimol injection. A localized visual RF was visible in the lower left visual field. The inset shows another RF from another channel along the injectrode shaft. The RF was slightly displaced, suggesting that the injectrode spanned multiple cortical columns in this session. **(E, F)** Similar pre-inactivation RF’s from two additional example sessions, this time from monkey A. **(G-I)** Saccade landing errors after muscimol injection for the same sessions as in **D**-**F**. For each session, there were increased landing errors for target locations corresponding to the pre-injection RF locations (dark blobs). Note that the scotomas were significantly larger than a single cortical column (the deficit in **G** matched the multiple RF locations of **D**), suggesting that our injected volumes were also large enough to inactivate all cortical layers (**B** and Methods). The scotomas were always larger than our presented visual stimuli (locations indicated schematically by green discs). Figure S1 shows additional scotoma maps from other sessions. The SC schematic in **A** was adapted from (107), and the numbers on the top-left of **G**-**I** and top-right of **B** are session identifiers.

Moreover, the SC itself projects back to the lateral geniculate nucleus (LGN) (22, 23) and other parts of the thalamus (24–32), to ultimately influence the cortex (4, 24, 26, 33–38). Even within the geniculate pathway itself, there are LGN projections to extrastriate cortex bypassing the primary visual cortex (V1) (39–50). Thus, a fundamental question in systems neuroscience is how such divergent anatomical pathways and loops can ultimately result in convergent function, with a purposeful and coherent influence on perception, cognition, and behavior.

In the case of primate visually-guided eye movements, a general consensus has been that retinotectal projections (5–7, 9–19, 25, 51) must be normally active and functional for behavior, not unlike in rodents (52–54). Evidence for this includes some sparing of visual responses in the SC after eliminating V1 activity (14), as well as the so-called phenomenon of blindsight after long-term V1 lesions (4, 9, 10, 12, 55–60), in which residual visual abilities sufficient to drive eye and arm movements are observed after enough post-lesion recovery and training have taken place (4, 12, 57, 61–72). However, it still remains possible that pathways other than the retinotectal one can contribute to mediating blindsight abilities (46, 47, 73–76), and these abilities also require substantial plasticity and reorganization (12, 32, 77). This leaves open the question of how the different visual pathways, including the retinotectal one, might operate in the normal brain.

Here, we approached the problem of understanding the roles of multiple visual-motor pathways by focusing on extremely short-latency ocular reflexes that are inevitable (78), unadaptable (79–83), and well-characterized in both humans (84–96) and monkeys (79–82, 97–99). We specifically studied the phenomenon called saccadic (or microsaccadic) inhibition, which refers to a short-latency reduction (and almost complete cessation) of saccade generation likelihood in response to a sensory transient (78). This motor resetting (98) phenomenon is essentially time-locked to short-latency visual responses (presumably in oculomotor structures like the SC) (81), but classic interpretations of its underlying mechanisms, such as via lateral inhibitory interactions in the SC or frontal eye fields (FEF) (85, 87, 89, 94, 100–103), are not experimentally supported (104–106).

We first performed acute V1 inactivation experiments and discovered that this was entirely sufficient to render the phenomenon of saccadic inhibition statistically unobservable. Then, using long term V1 lesions, we found that latent visual signals bypassing the geniculostriate pathway must still exist, which, combined with computational models, then enabled us to identify evidence for such latent visual signals even in our acute V1 inactivation experiments, albeit not directly on the inhibition phenomenon itself. Besides aiding in identifying the pathway origins of a well-known ocular reflex, our results help clarify and reconcile conflicting views and assumptions about the roles of different retinal projections in primate visually-guided motor behavior.

## Results

### Reversible V1 inactivation abolishes saccadic inhibition

Our goal was to investigate the relative contribution of the geniculostriate pathway, versus the retinotectal (3–5, 9–16, 57) or other (4, 20, 21, 35, 39–47, 56, 57) pathways, in visually-driven saccadic inhibition. To do so, we first reversibly inactivated a small portion of the primary visual cortex using muscimol injection (Methods; Fig. 1A), and we confirmed the effectiveness of cortical inactivation by simultaneously recording multi-unit activity along the injectrode shaft during the injection period (Fig. 1B). Across sessions in two monkeys, we targeted extrafoveal regions in either the right or left visual field representation of V1 (Fig. 1C).

We confirmed the affected visual field regions using different approaches. Prior to injection, but with the injectrode partially in the cortex, we recorded multi-unit activity and mapped the visual receptive fields (RF’s) that were neighboring our penetrated injectrode site (Methods). Example RF’s from one session in each visual hemifield that we tested in each monkey are presented in Fig. 1D-F. In all cases, expected V1 RF’s were encountered. After the injection period (Methods), and after multi-unit activity confirmation of cortical inactivation (Fig. 1B), we ran a modified version of a classic visually-guided saccade task, in which the monkey received instruction to look at an eccentric target by removal of the central fixation spot. This central fixation spot was always foveally visible even during cortical inactivation (Methods). Across trials, the saccade target could appear at any location on the display (most of which were visible to the monkey because of the focal nature of our injection). However, on some random trials, we presented the target at a location where we expected a visual field scotoma to exist (based on RF’s like those of Fig. 1D-F). We then measured the saccade landing errors. Critically, we did not penalize the monkeys for having a large error, and we never repeated an error trial so that the monkeys did not learn to make a memory-guided saccade to a specific region of the display. Consistent with the development of a cortical scotoma, landing errors were markedly increased in visual field locations around the RF locations recorded prior to the injection (compare the behavioral deficits – dark blobs – in Fig. 1G-I to the RF locations in Fig. 1D-F). In fact, the spread of the behavioral scotoma on a given session still reflected the pre-injection electrode contact locations. For example, the inset in Fig. 1D shows a second RF encountered at another depth of the injectrode from the same session. This RF was slightly displaced in location from the first one, suggesting that the penetration was not directly parallel to a single cortical column; the visual field deficit in Fig. 1G reflected the aggregate locations of the two RF regions. Thus, we confirmed that in each session, there was a confined region of the visual field in which each monkey was rendered cortically blind.

After confirming cortical inactivation, we ran a fixation task in which a visual onset (black disc of 0.18 deg radius and 100% Weber contrast relative to the background) could appear either in the affected visual field location, or in the opposite, unaffected hemifield (at the same vertical position, but with a reflected horizontal position). We also interleaved no-stimulus trials, in order to assess saccade rates in the absence of any stimulus-driven saccadic inhibition. Cortical inactivation had a dramatic influence on the inhibition phenomenon, and this can be seen from the raw saccade onset raster plots of Fig. 2. In these plots, each row is an individual trial, and each dot indicates the onset time of a fixational saccade. For each monkey, when the stimulus onset appeared with an intact V1, there was a short-latency, reflexive inhibition of saccade generation (78, 79, 89, 101). This reflex was so robust that it persisted across thousands of trial repetitions in each monkey (greenish dots in Fig. 2), consistent with prior knowledge (79–83), and justifying our pooling of the data across sessions in this and most subsequent analyses (Methods). When the same stimulus appeared in a region of the visual field for which V1 activity was eliminated, no saccadic inhibition was visible, and the saccade rasters did not abruptly change after stimulus onset (reddish dots in Fig. 2). This was despite the fact that retinotectal and other sensory-motor pathways (Fig. 1A) were still fully intact, and thus possessed putative visual responses that could, at least theoretically, reach the oculomotor control system and contribute to saccadic inhibition.

**Figure 2.**
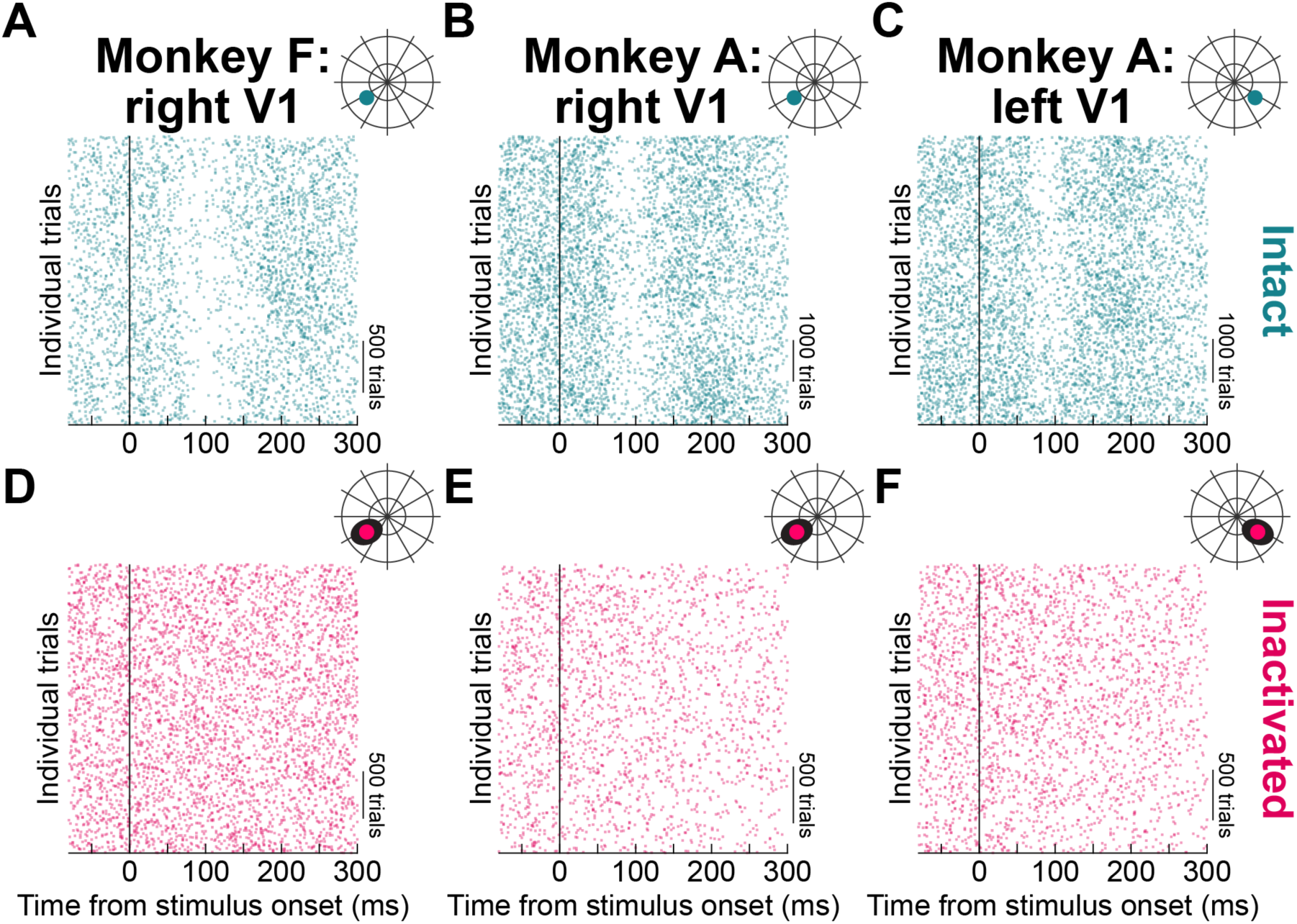
V**1 inactivation abolishes saccadic inhibition. (A)** Fixational saccade onset time rasters, as a function of time from visual stimulus onset (x-axis) and trial number (y-axis), across all of the intact V1 sessions of monkey F. A robust inhibition of fixational saccade generation occurred with a short latency of <100 ms from stimulus onset (79–81, 98, 104, 109). **(B, C)** Similar observations from monkey A. The visual field schematics shown above each panel indicate the approximate locations of the appearing visual stimuli in all sessions (also see Fig. 1C). **(D-F)** When V1 was inactivated, saccadic inhibition disappeared, as evidenced by the unaltered density of saccade onset times shortly after stimulus onset, and this happened for both right (**D**, **E**) or left (**F**) V1 inactivation. The visual field schematics above each panel show the approximate scotoma regions (black ovals) and the visual stimulus locations (reddish discs). Also see Fig. 3 for quantification of the inactivation effects.

Quantitatively, we confirmed two key observations about the effects of transient V1 inactivation on stimulus-driven saccadic inhibition, which we describe next. First, there was a clear difference between saccade rates on inactivation and intact trials, for the very same stimulus locations; this reveals, by far, the largest effect on the stimulus-driven saccade rate signature from all previous perturbation experiments in other brain areas investigating it (105, 106, 108). Second, there was no difference between saccade rates on inactivation and no-stimulus trials; this means that V1 inactivation rendered stimulus-driven saccadic inhibition statistically unobservable.

To demonstrate the first key observation, we plotted saccade rate as a function of time from stimulus onset from the intact and inactivation trials (Fig. 3A-C); the two data sets show matched stimulus locations (per pair of inactivation and intact sessions) for each monkey, with the only difference being whether the trials were collected with an intact (greenish curves) or inactivated (reddish curves) V1. With an intact V1, saccade rate inhibition followed by rebound took place (80, 81, 89, 91, 92, 98), and this resulted in two post-stimulus epochs during which the saccade rate was statistically significantly different from that in the inactivation trials: the first epoch is indicated by blue on the x-axis in Fig. 3A-C, and shows that the rate on intact trials was first lower than the rate on inactivation trials; the second epoch is indicated in red on the x-axis in Fig. 3A-C, and shows that the rate on intact trials was higher than on inactivation trials (all statistical results are reported in Table S1). This dichotomous difference from inactivation is consistent with either a weakened, or completely absent, stimulus-driven saccade rate modulation during the inactivated trials.

**Figure 3.**
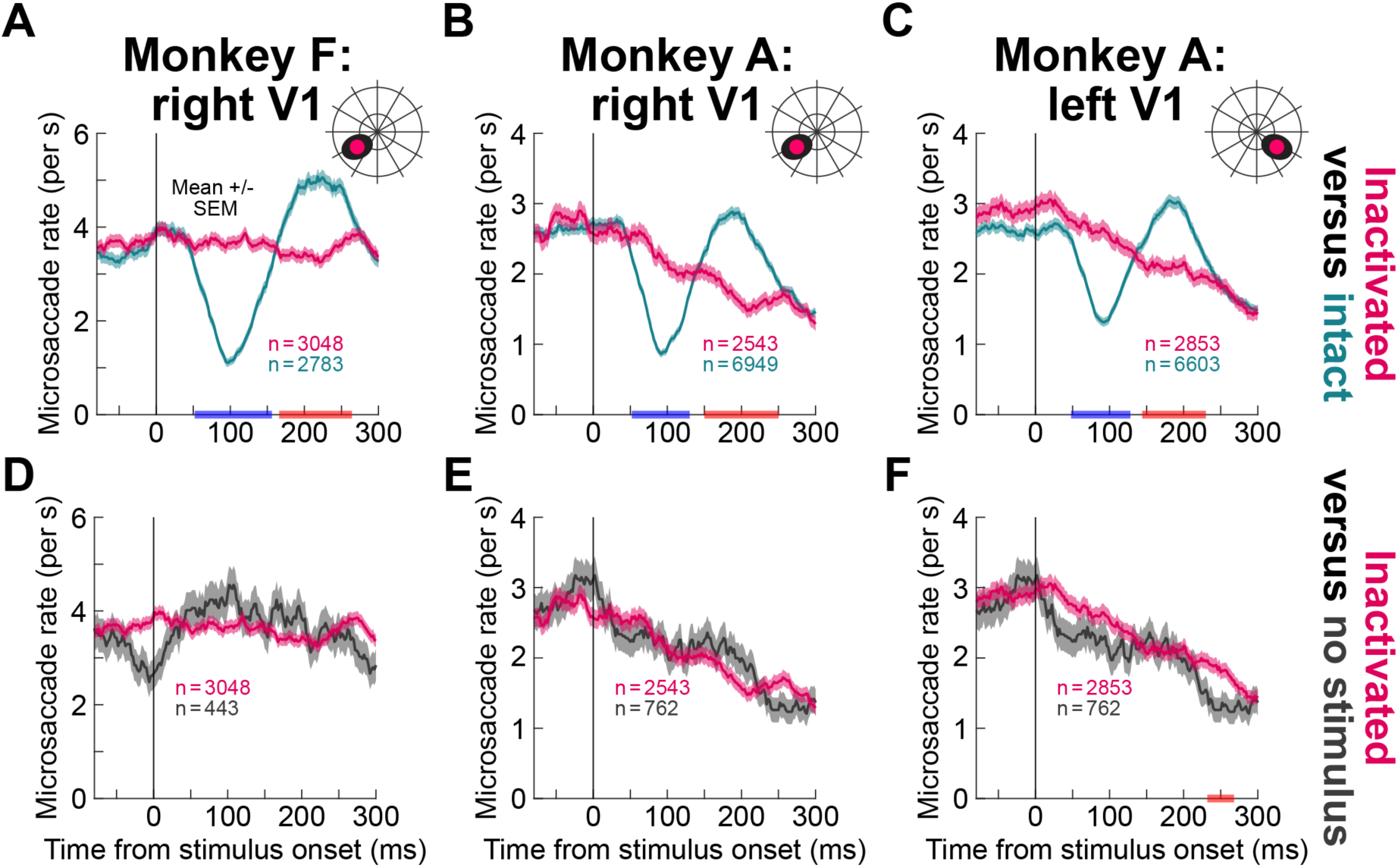
Saccade rates during V1 inactivation are indistinguishable from no-stimulus saccade rates. **(A)** Same data as in Fig. 2A, but expressed as fixational saccade (microsaccade) rates (Methods). With an intact V1, short-latency saccadic inhibition followed by rebound was observed; with inactivated V1, the saccade rate was not modulated by stimulus onset when the stimulus was presented within the V1 scotoma (inset schematic above the figure). The blue and red markings on the x-axis indicate the epochs during which the two curves differed from each other, based on cluster-based statistical comparisons (Methods and Table S1). **(B, C)** Similar to **A** but for monkey A’s data in either the right (**B**) or left (**C**) V1. Note that this monkey had a time-varying saccade rate trend even with an intact V1 (for example, the rate in the intact curves at 300 ms was lower than the pre-stimulus rate). Such a time-varying rate was also evident in this monkey without any stimulus as seen in **E**, **F**. **(D-F)** To check whether the results of **A**-**C** reflected merely a weakened saccadic inhibition effect, as opposed to a statistically eliminated one, we plotted the inactivation data of **A**-**C** (reddish curves) together with no-stimulus data (gray curves). During saccadic inhibition epochs (starting <100 ms after stimulus onset; **A**-**C**), there was no difference between saccade rates with V1 inactivation and saccade rates with no stimulus (Methods; also see Fig. S2). For monkey A and left V1 (**F**), there was a late epoch showing a difference, but this was >100 ms after any saccadic inhibition was expected to occur, and it showed that inactivation rate was slightly higher, not lower, than the no stimulus rate. Error bars denote SEM across trials.

To decide between whether the inactivation effect was merely weakened versus absent, we then compared the inactivation saccade rates to those obtained without any visual stimulus at all. We found no significant differences between the two situations, especially during the critical time epochs of saccadic inhibition (within <100 ms from stimulus onset). These results are shown in Fig. 3D-F, and they confirm that the time-varying saccade rates observed in Fig. 3A-C during V1 inactivation (particularly for monkey A) were not stimulus-driven; they likely reflected other factors related to expectations on the timings of successive trials, and they were still present in the no-stimulus trials. Naturally, we also confirmed that the intact V1 data still showed the dichotomous difference from no-stimulus trials expected from the data of Fig. 3 (Fig. S2A-C), and we additionally established that no-stimulus trials showed similar saccade rates whether V1 was intact or inactivated (Fig. S2D, E). This indicates that the intrinsic ability of the monkeys to generate saccades, in general, was not altered by V1 inactivation; only the saccadic resetting process itself was abolished. Finally, the results of Fig. 3A-C were also visible in individual sessions (Fig. S3).

Our task also had a built-in control condition, in that we interleaved trials during which the stimulus appeared in the opposite hemifield to the one targeted with muscimol. Here again, saccadic inhibition was normal, as might be expected from the fact that the stimulus now activated unaffected V1 neurons. This can be seen in Fig. S4A-C. Saccadic inhibition was unaltered in all cases, but the post-inhibition saccade rebound, which is a cortical phenomenon (80, 104, 106, 110–112), was sometimes modified. Thus, our results so far demonstrate that reflexive saccadic inhibition necessarily depends on the occurrence of stimulus-driven neuronal activity in V1; if that activity is absent, no stimulus-driven saccadic inhibition takes place.

We had also previously shown that the detailed properties of saccadic inhibition, but not the phenomenon as a whole, depend on stimulus luminance polarity (80). Given that V1 has a majority of neurons preferring dark stimuli (113–115), which may or may not be reflected in the alternative visual pathways bypassing V1 (116), we next wondered whether our effects depended on luminance polarity. To test this, we presented bright discs, again with 100% contrast (Methods). Once again, there was a dramatic effect of V1 inactivation on the stimulus-driven saccade rate signature, and saccade rates were not different from those without a stimulus onset (Fig. S5).

In a similar vein, saccadic inhibition exhibits feature tuning properties to visual stimulus parameters like contrast (81, 89, 117–119). Therefore, we also tried stimuli of different Weber contrasts (10% versus 100%). As expected (81, 89), in the intact case, saccadic inhibition was rendered weaker and later than with 100% contrast stimuli. However, it was still present, and it was still abolished with V1 inactivation (Fig. S6).

Thus, the effect of V1 inactivation is to cause an all-or-none change in the occurrence of saccadic inhibition: as long as a stimulus, however weak, can elicit V1 responses, saccadic inhibition can still be observed; otherwise, saccadic inhibition is eliminated.

### V1 necessity does not depend on task context

In the experiments described so far, the monkeys were fixating a central spot, but did not have to otherwise react to the eccentric stimulus. Might it be the case that engagement in a more goal-directed behavioral response was a necessary prerequisite for alternative visual pathways to be functionally activated? To test this, we resorted to a cognitively-demanding memory-guided saccade paradigm. Here, each monkey was expected to hold the position of a briefly flashed stimulus in memory while maintaining fixation. Then, when the fixation spot was eventually removed, the monkey had to generate a memory-guided eye movement towards the remembered location (Methods). Since the monkey was cortically blind on the injection trials, we interleaved trials in the opposite hemifield so that the monkey could infer the task context. This was no problem, and both monkeys readily performed the paradigm.

Their landing errors in the intact hemifield were, naturally, larger than those in the visible trials of Fig. 1G-I (outside the dark scotomas of that figure), and they were closer in absolute value to those in the inactivated hemifield (Fig. S7). This confirms that the monkeys were relying on memory in both cases, and that they explicitly paid attention to the inactivated region during the trials in which they did not consciously perceive the brief flash at trial onset.

Despite the strong task engagement, saccadic inhibition after brief flash onset was again eliminated with V1 inactivation. This can be seen in Fig. 4, with the data being similarly formatted to Fig. 3. With an intact V1, brief flashed stimuli at trial onsets elicited robust saccadic inhibition, which was gone with V1 inactivation (Fig. 4A, B). Moreover, saccade rate was no different from that on no-stimulus trials (Fig. 4C, D), but saccadic inhibition was intact when the flashes appeared outside of the visual field region affected by V1 inactivation (Fig. S8).

**Figure 4.**
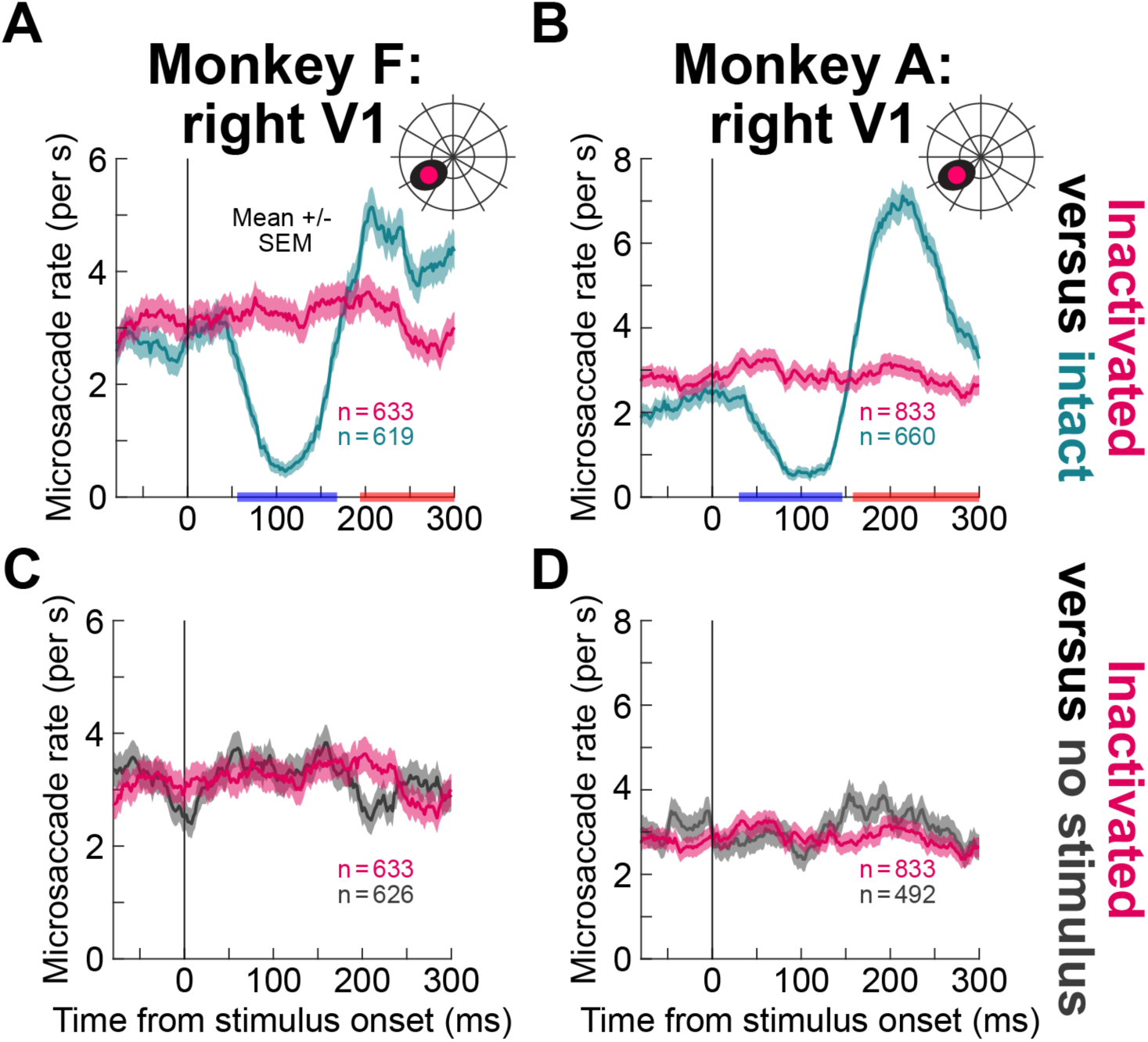
Similar results with a cognitively demanding task. (A, **B)** Like in Fig. 3A-C, V1 inactivation eliminated saccadic inhibition even when the task required attention to a particular visual field quadrant (Methods). In these panels, we compared fixational saccade (microsaccade) rates with and without inactivation, revealing the same dichotomous difference from the intact V1 data as in Fig. 3A-C. **(C, D)** We then compared the inactivation data to no-stimulus data. There was no difference between the reddish curve and the dark gray curve, just like in Fig. 3D-F. Thus, the results of Fig. 2, 3 held even in the context of a different behavioral task. Error bars denote SEM across trials, and all other conventions are like in Fig. 3.

We also repeated all experiments above while injecting saline, rather than muscimol, in V1 of each monkey. Saccadic inhibition was not affected by these injections, suggesting that the muscimol effects were not due to volume displacement of neuronal tissue by the injected fluids (Figs. S9, S10A, B, E, F).

Therefore, visually-driven reflexive resetting of the saccadic system (78) is an exclusively cortical phenomenon, which does not depend on either the feature properties of the appearing stimuli or the behavioral task context.

### Permanent V1 loss unmasks a role for alternative visual pathways

Because the above results are contrary to anatomical (3, 4, 6–8, 10, 11, 15, 16, 19, 57, 120) and physiological (12, 14, 121–124) evidence of visual sensory pathways bypassing V1, we wondered whether and to what extent latent visual signals in these alternative visual pathways can still be manifested. We resorted to three additional approaches, one experimental, one theoretical, and one analytical. In what follows, we describe these approaches in order.

In the experimental approach, we used permanent, rather than transient, V1 inactivation (77, 125, 126). In monkey models of V1 lesions, after some post-lesion recovery time and training, the monkeys exhibit residual visual and cognitive abilities (9, 12, 56, 61–63, 68, 71, 127–134) that allow orienting towards stimuli in the blind hemifield (12, 61–64). This monkey-equivalent to so-called human blindsight (4, 10, 55, 58–60) suggests that there is an available substrate of visual information bypassing the geniculostriate pathway. Therefore, we decided to check whether saccadic inhibition occurs in monkeys with permanent V1 lesions.

In three monkeys, we measured fixational saccade rates around stimulus onset well after the monkeys developed blindsight-like abilities (Methods). For example, for monkeys Tb and Ak, evidence of their residual visual abilities was already published (62, 63, 130, 135). Two out of the three monkeys showed clear stimulus-driven saccadic inhibition when a visual target appeared in the hemifield represented by the permanent V1 lesion (Methods). This can be seen in Fig. 5A-C. Monkey Tb showed almost normal inhibition followed by rebound (Fig. 5A), and so did monkey S (Fig. 5C); the differences in time courses between the two monkeys are likely due to significant differences in the stimulus conditions (Methods). However, in monkey Ak, we did not find such evidence (Fig. 5B). This could either mean that recovery of saccadic inhibition is not always possible, or it could also be due to the noisy nature of this monkey’s eye data, as well as the monkey’s low saccade rates (Methods).

**Figure 5.**
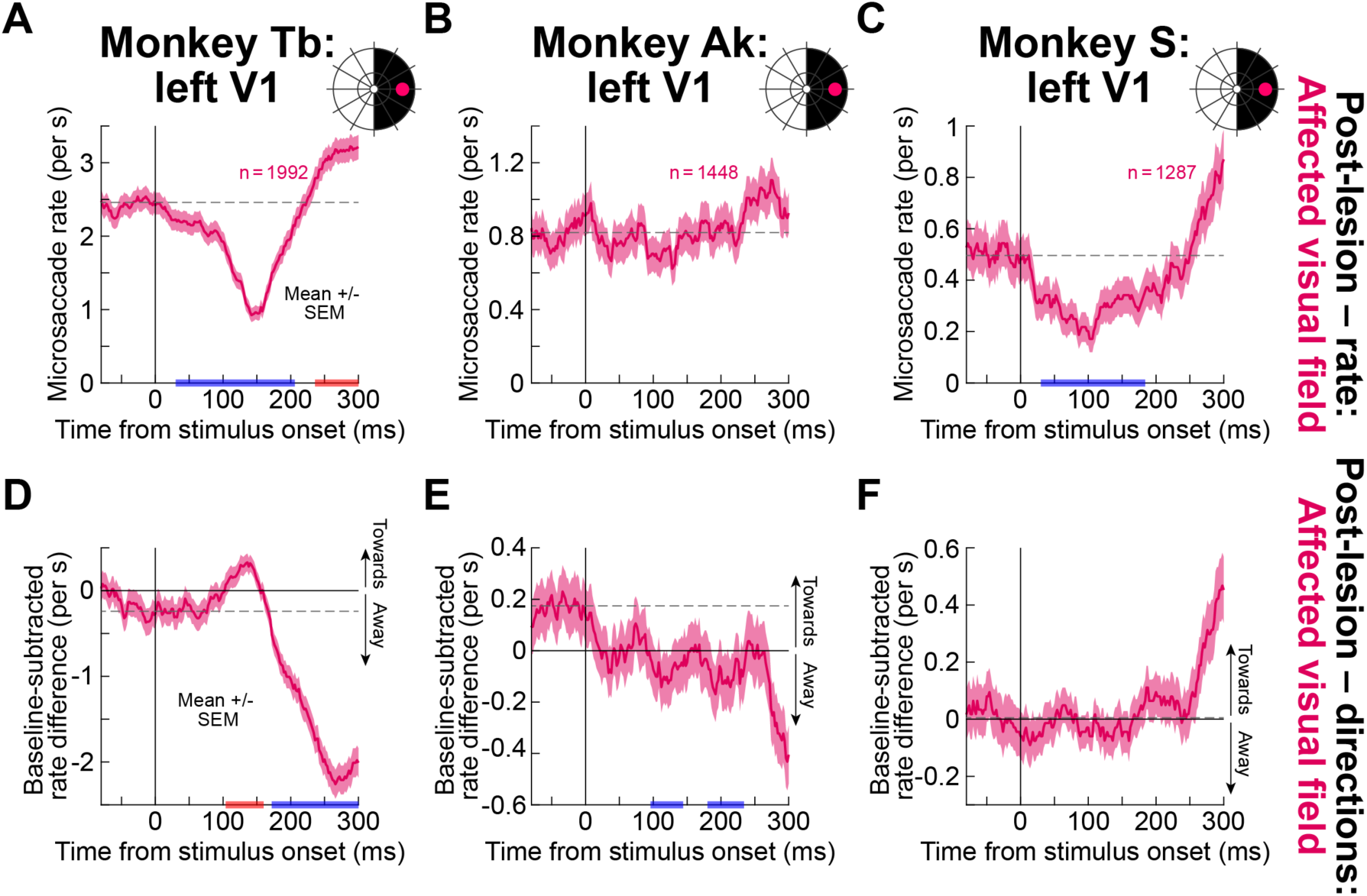
Saccadic inhibition can be observed with permanent V1 lesions. **(A)** In a monkey living with a permanent lesion of extrafoveal left V1, presentation of a visual stimulus in the affected hemifield (schematized as black in the inset above the figure) resulted in saccadic inhibition followed by post-inhibition rebound (blue and red x-axis epochs describe statistically significant differences from the pre-stimulus baseline), like with an intact V1. **(B)** This was not the case in a second monkey from the same experiment. **(C)** However, a third monkey in a different experimental setting (Methods) did exhibit saccadic inhibition after the V1 lesion. **(D)** For the same data as in **A**, we plotted the baseline-subtracted difference between fixational saccade rate towards and away from the visual stimulus location (82) (Methods), allowing us to explore direction biases associated with early saccadic inhibition phases (90, 91). Positive deflections mean biases in saccade directions towards the hemifield of the appearing stimulus. Blue and red epochs on the x-axis indicate when the directions were statistically significantly different from the pre-stimulus directional distribution (Methods and Table S1). During saccadic inhibition in **A**, the fixational saccade directions were biased towards the visual stimulus location, as expected in intact animals (79, 82, 88, 90–92, 98). There was later a reversal opposite the stimulus direction. **(E)** When there was no saccadic inhibition observed, the direction oscillation was less clear and diffuse in time. **(F)** There was also no clear direction oscillation in the third monkey. Error bars denote SEM.

Interestingly, when we looked at saccade rates for stimuli in the intact visual field, we also found weak (or non-existent) initial saccadic inhibition in this monkey, but similar later modulations to monkey Tb (Fig. S11). However, this was still altered relative to pre-lesion behavior: monkey Ak clearly had saccadic inhibition in that case (Fig. S12). Thus, in at least two out of the three monkeys (Fig. 5A, C), saccadic inhibition could still be driven by alternative visual pathways after permanent V1 loss. A latent visual signal in the alternative visual pathways does exist.

We also analyzed the eye movement directions in the three lesion monkeys, and we did so using the same technique that we recently described (82) (Methods). Our motivation for analyzing directions here was that in our type of task, the fixational saccades that do happen around the time of saccadic inhibition become also biased in directions (90, 91). While it has been known for more than two decades that such directional modulations exist around the time of saccadic inhibition (79, 88, 90–92, 98, 136, 137), recent theoretical and experimental accounts suggest that the modulations can more directly reflect visual signals in spatial topographic maps, such as in the SC, which are part of the alternative visual pathways (e.g. retinotectal projections) (78, 82, 104, 105). Moreover, these theoretical accounts suggest that the neuronal origins of direction modulations might be anatomically distinct from those associated with temporal inhibition (78, 104). Once again, in monkey Tb, there were clear saccade direction oscillations after permanent V1 loss, with early saccades around the time of rate inhibition being biased towards the direction of the stimulus presented in the lesioned visual field, and later saccades being biased in the opposite direction (Fig. 5D). This is the clearest evidence from these lesion studies that a latent visual signal bypassing V1 Does exist. In monkey Ak, there was a less clear directional modulation, with later saccades being biased opposite to the stimulus location (Fig. 5E). Given that there was no temporal resetting in the rate curve (Fig. 5B), it is expected that directional effects might be diffuse in time in this case. There was also no clear bias towards the stimulus location in the affected hemifield for monkey S (Fig. 5F). In all three monkeys, when there was clear inhibition either in the intact hemifield (Fig. S11) or in pre-lesion tests (Fig. S12), more consistent stimulus-driven saccade direction modulations occurred. Therefore, saccade directions in lesion monkeys added further evidence that a latent visual signal bypassing V1 might still be relevant, at least under the right conditions. This motivated us to proceed to the next theoretical and analytical steps alluded to above.

### Weakened visual drive can impair saccade rate more than direction effects

Motivated by the lesion results, we then asked a more theoretical question regarding our transient inactivation observations of Figs. 1-4: is it at all possible for saccade direction effects to be observable with weakened visual inputs, even when saccade rate inhibition is rendered statistically unobservable? We developed a computational model of saccadic inhibition, in which we could titrate the strength of the visual signal causing temporal resetting. The intact model, an even simpler version of one we presented earlier (98, 138), demonstrates how exogenous visual inputs rapidly achieve two independent outcomes: they countermand saccade generation plans to implement phase resetting (saccadic inhibition), and they also transiently bias spatially organized topographic circuits (via localized visual bursts) to modulate saccade directions. This model captures all salient aspects of saccadic inhibition from experiments, including saccade direction modulations, and this can be seen from an example intact model simulation of 2000 trials (Fig. 6A, B).

**Figure 6.**
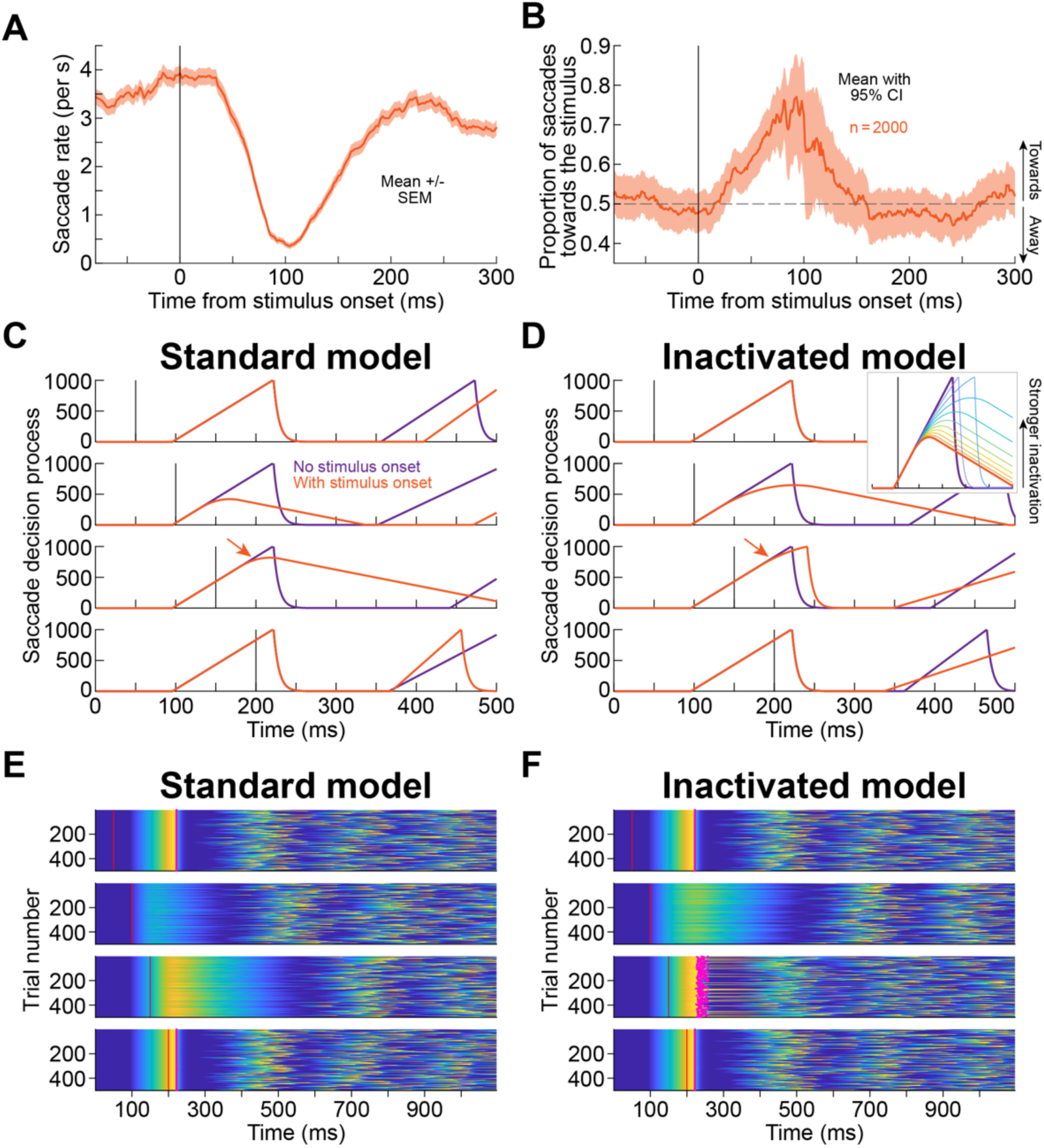
Saccadic inhibition as a visually-driven countermanding of endogenous saccade plans. **(A)** Saccade rate from 2000 simulated trials of our computational model (Methods). Error bars denote SEM across trials. **(B)** Proportion of saccades in the direction of the stimulus as a function of time from stimulus onset, for the same data as in **A**. Error bars denote 95% confidence intervals. The model replicated both rate (**A**) and direction (**B**) modulations after stimulus onset. **(C)** In the model, a stimulus onset (black vertical line in each row) countermands an endogenous saccade plan by slowing down, and reversing, a rise-to-threshold process. Each row shows an initial endogenous saccade plan (dark purple) and the same plan if a stimulus onset occurs (orange). If the stimulus appears before the saccade process starts, the saccade proceeds unimpeded (first row). If the stimulus appears early enough in the saccade plan (second and third rows), the countermanding succeeds, and a subsequent saccade process is later initiated. If the stimulus appears late in the plan, the countermanding fails, and the original saccade is triggered anyway (“escape” movements). **(D)** With V1 inactivation, the efficacy of the visual signal in countermanding is impaired. Here, we simulated a scaling down of the visual signal by 60% for illustration. Now, the saccade in the third row fails to be countermanded and is triggered nonetheless (red arrow). Thus, overall, there is less delaying of saccades. The inset shows how different scalings of the visual input’s impact (between 0% and 100%, in steps of 10%) alter the countermanding process. **(E, F)** Repeated simulations of scenarios like in **C**, **D**, demonstrating how the timing of subsequent saccades can be affected by stimulus onset. Here, we kept the initial saccade the same but ran the stochastic model for all subsequent saccades. Each row is a saccade trial, and the color indicates the value of the rise-to-threshold process. If countermanding succeeds, subsequent saccades are delayed (second and third rows in **E**). If it fails, the first saccade is triggered (magenta dots indicate the threshold crossing of the accumulator on each trial). A weakened visual signal impairs countermanding and reduces the delaying (and thus the inhibition; third row in **F**).

To briefly explain the mechanistic insight of the model in the intact case, saccadic inhibition occurs in the model because a visual input starts a countermanding process attempting to cancel an ensuing saccade plan; this is achieved via altering the rate of a rise-to-threshold accumulator process for saccade generation (which is physiologically equivalent to raising the threshold for saccade triggering (78)). If the visual input comes early enough in the rise-to-threshold process, the change in accumulator rate eventually causes the saccade to get completely canceled (Fig. 6C; second and third rows); if the visual input is late, the rate alteration of the accumulator is too slow, and the threshold for triggering the planned eye movement gets reached nonetheless (Fig. 6C; fourth row; Methods). Across repetitions, saccade likelihood will be altered by stimulus onset, as a simple side effect of delaying and/or canceling saccade plans (Fig. 6E). For saccade directions, the model assumes that any given saccade has a tendency to be directed opposite the previous saccade; this is an experimental reality, especially when fixating a small spot (90, 98, 109, 138–140). Once the visual stimulus is processed, a single eye movement plan, the current one at the time of stimulus processing completion, is biased in the direction of the stimulus; this reflects visual bursts in spatially-organized maps like in the SC (141), and all subsequent eye movements revert to obeying the directional rule of steady-state saccade generation.

To simulate the effect of transient V1 inactivation in the model, while still maintaining a putative latent visual signal (e.g. in alternative pathways), we weakened the impact of the visual input on the rate of the rise-to-threshold process by a scale factor (called visual sparing factor; Methods). For example, for the same initial eye movement plan of Fig. 6C, E, if the efficacy of the countermanding process by exogenous stimulus onset is reduced by 60% (i.e. 40% visual sparing factor), then some stimulus onsets (e.g. at 150 ms; third row in Fig. 6D) fail to successfully countermand the saccade plan (the inset shows how successively weaker latent visual signals impair the countermanding efficacy); the net result is that across many repetitions, subsequent saccades are less “delayed” by exogenous stimulus onsets (Fig. 6F; compare the third row to that in Fig. 6E), resulting in weakened saccadic inhibition.

Critically, according to the model, it is perfectly plausible for saccadic inhibition to be statistically eliminated when there is still a non-zero latent visual signal present. Consider, for example the results shown in Fig. 7A, B. Here, we ran the model with a weakened visual input’s impact by a factor of 90% (mimicking an approximate 10% sparing of visual inputs from alternative pathways if the primary geniculostriate one is lost (4, 7, 16)). In this simulation, we ran 2000 model trials, and then characterized saccade rate and directions around stimulus onset, like in the intact case above (the intact model curves are replicated from Fig. 6A, B for easier comparison). The rate modulation was rendered absent, but a stimulus-driven biasing of saccade directions, while weaker than with a fully intact model, was still present. This indicates that a latent visual signal bypassing the geniculostriate pathway can still be observable in saccade directions even if saccade rate inhibition is not (and this can happen without the need for long-term recovery from permanent lesions). This idea is further corroborated by Fig. 7C, D in which we simulated a no-stimulus case; here, the direction effects were eliminated as might be expected (note that the rate fluctuation seen in Fig. 7C at around 100 ms is a result of numerical simulations with several random number generators for the different model parameters). Thus, weak latent visual signals can still be seen in saccade direction biases even when rate inhibition is eliminated.

**Figure 7.**
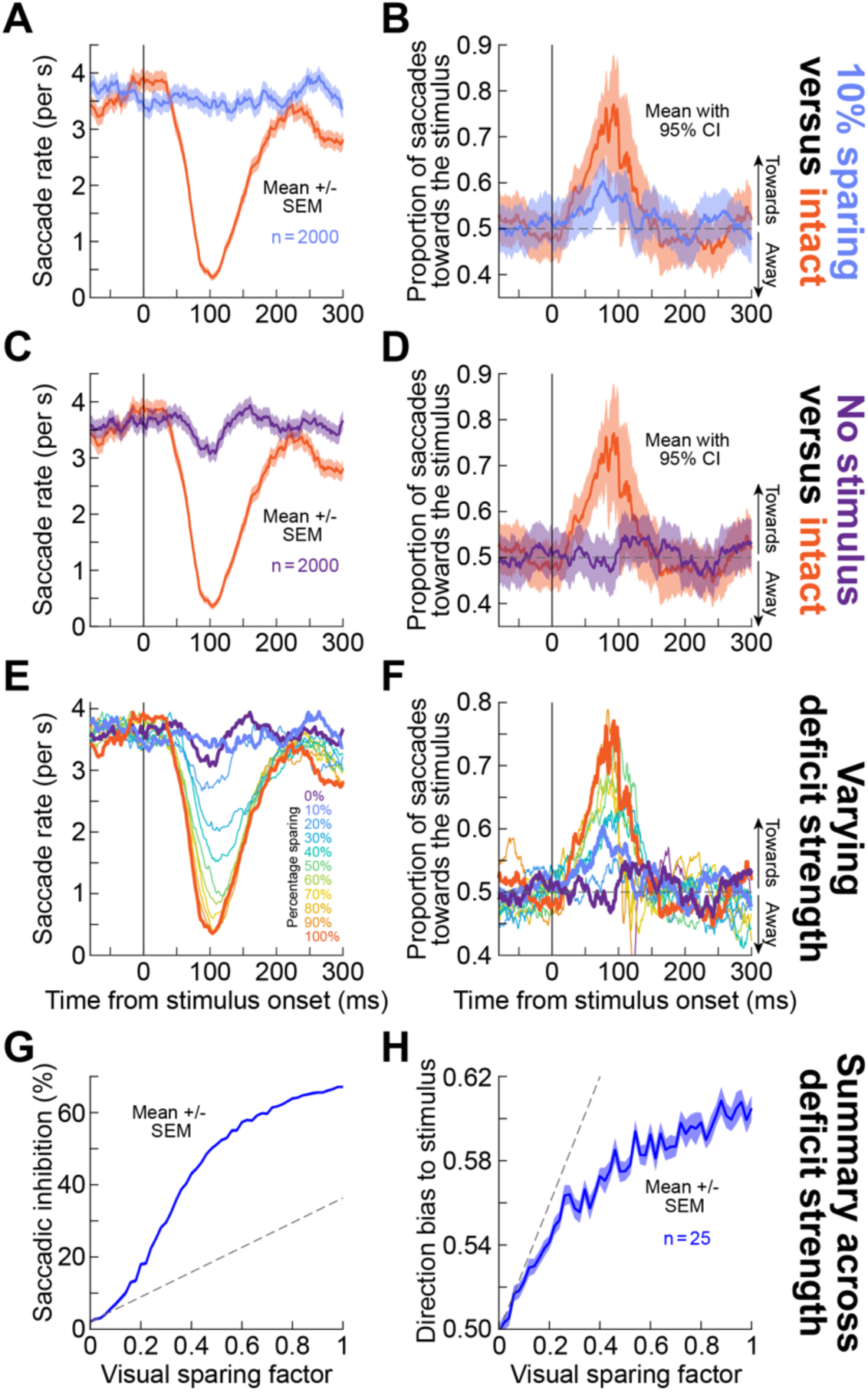
A latent visual signal can influence saccade directions even when rate inhibition is statistically unobservable. (A, **B)** The orange curves are like in Fig. 6A, B (intact model). In light purple, we ran an “inactivated model” with a sparing of 10% latent visual signal influencing saccadic countermanding (Methods). Saccadic inhibition was eliminated (**A**), but there was still a directional biasing (**B**). Error bars denote SEM in **A** and 95% confidence intervals in **B**. **(C, D)** When we ran the model with no stimulus (0% sparing of a visual signal), both the rate and direction modulations were eliminated; note that there was an apparent rate oscillation at around 100 ms that is due to the stochastic nature of the model simulations (Methods). **(E, F)** For different amounts of latent visual signal sparing, there were systematic alterations in saccadic inhibition and direction modulations, as might be expected. **(G, H)** Critically, weak visual signals were more effectively measurable in the model’s direction rather than rate modulations. For each latent visual signal strength, we ran 25 instantiations of the model (each simulating 2000 model trials), and we then characterized the amount of saccadic inhibition (**G**) and the amount of stimulus-driven direction modulation (**H**; Methods). At weak visual signals, the rate effect was at a floor level and the curve started with a lower slope than at high latent visual signal strengths (the tangent line in **G** was below the curve). On the other hand, the direction effect curve started with a high slope already at the lowest visual signal impacts (**H**; tangent line was parallel to or above the curve), suggesting that the latent visual signal could still influence saccade directions when it was too weak to cause measurable saccadic inhibition. The tangent line slope was calculated based on sampling the y-axis value at the two points at 0.06 and 0 countermanding efficacy on the x-axis. Error bars here denote SEM.

We also confirmed the robustness of this observation by running different instantiations of the model at different levels of visual signal sparing. In each instantiation, we ran 2000 model trials, and we characterized saccade rate and directions. Weakening the visual input weakened both the inhibition and the direction oscillations (Fig. 7E, F). Then, we ran 25 instantiations at each visual sparing factor level (each instantiation simulating 2000 trials) and characterized either inhibition strength or direction modulation strength (Methods). At the weakest visual signal strengths (up to around 10% sparing factor), rate effects reached a floor effect (indicating elimination of saccadic inhibition) (Fig. 7G), but direction effects were still observable and scaled proportionally with increasing visual input strength (Fig. 7H; compare the slopes of the tangent lines in Fig. 7G, H). Therefore, even with a transient inactivation (like in Figs. 1-4), it is theoretically possible to lose saccadic inhibition as a “rate” phenomenon but still observe some directional influences of weak latent visual signals bypassing the geniculostriate pathway. With this new insight, we revisited our muscimol inactivation data, now focusing on saccade direction modulations.

### Direction modulations with transient inactivation suggest a weak latent visual signal bypassing V1

If the above lesion and model predictions are true, then we should have seen subtle stimulus-driven fixational saccade directional biases during transient V1 inactivation as well. Thus, our third and final (analytical) investigation of potential latent visual signals involved analyzing direction modulations in our original muscimol experiments. In the intact case, each monkey showed strong directional oscillations (Fig. 8A-C), as expected (79, 82, 88, 90–92, 98). Note that monkey A had a strong pre-stimulus directional bias that we characterized in great detail recently (82), and that created a dependence of the starting phase of the directional oscillation on the visual hemifield of the appearing stimulus. Thus, for this monkey, the modulations in the right and left visual fields were presented separately, also to reflect the fact that we injected muscimol in either the left or right V1 of this monkey.

**Figure 8.**
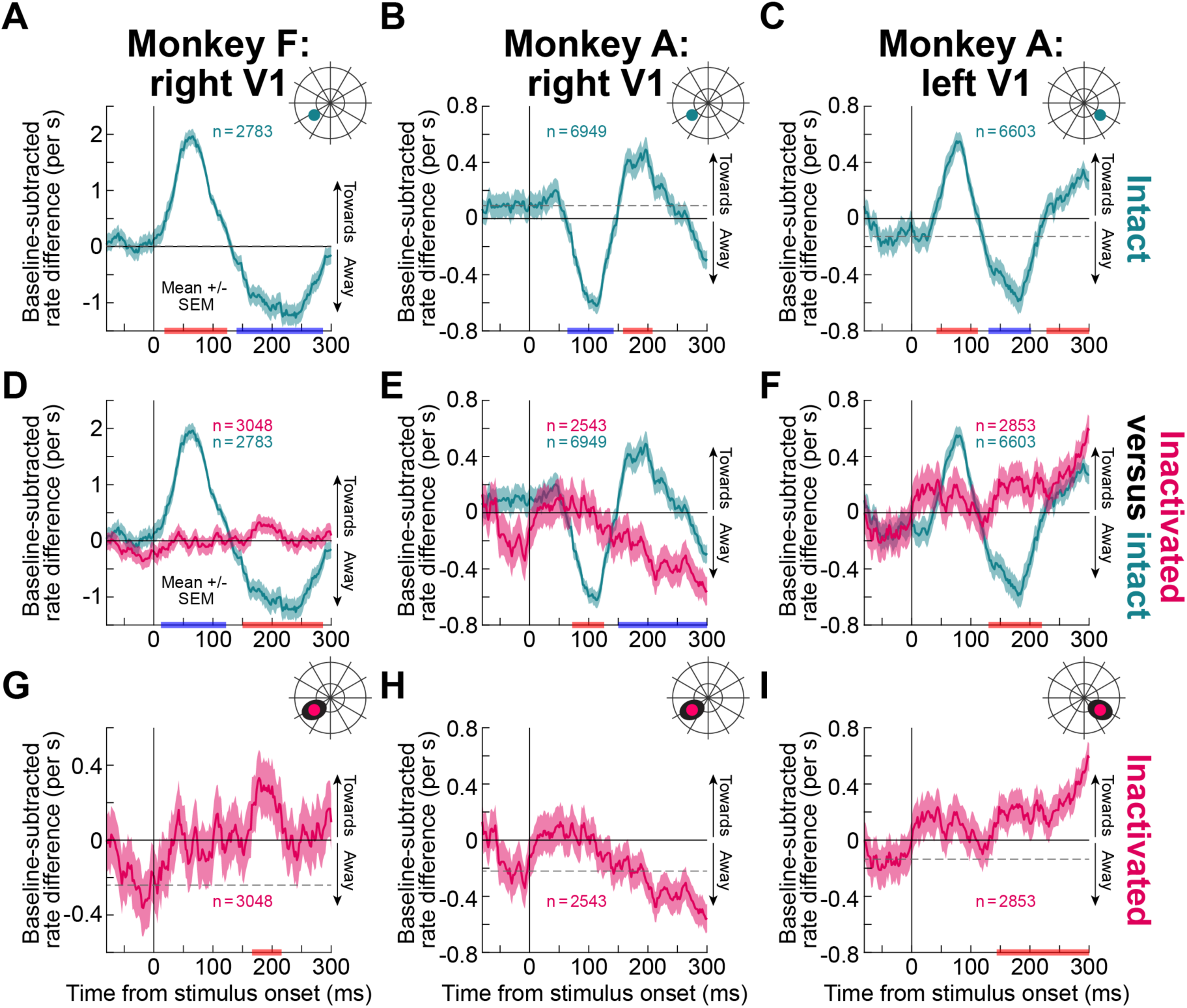
Evidence of a latent visual signal in fixational saccade direction biases after muscimol V1 inactivation. (A-C) Without V1 inactivation, each monkey showed strong fixational saccade direction oscillations after stimulus onset (shown here are the results for the stimuli that would appear in the scotoma region after muscimol injection). The figure is formatted similarly to Fig. 5D-F. **(D-F)** With reversible V1 inactivation (reddish curves), the direction oscillations were strongly altered relative to the intact situation (greenish curves; replicated from **A**-**C** for easier comparison), consistent with the elimination of saccadic inhibition in Figs. 2, 3. Blue and red epochs on the x-axes indicate significant differences between the conditions (Table S1). **(G-I)** Nonetheless, weak biases in saccade directions (relative to pre-stimulus baselines; dashed horizontal lines) were still visible, consistent with the modeling results of Figs. 6, 7. Here, the same reddish curves of **D**-**F** are plotted but with higher magnification and statistical comparisons to pre-stimulus baselines. We saw weak directional biases that were always in the same direction as the intrinsic short-latency direction modulation bias of each monkey and each hemifield without V1 inactivation (the first phase of the direction oscillations in **A**-**C**). Thus, despite the loss of saccadic inhibition, readout of spatial maps by saccadic eye movements reflected a very weak, latent visual signal bypassing the geniculostriate pathway. Error bars denote SEM. All other conventions are similar to Figs. 3, 5.

Critically, when V1 was inactivated, there were still very weak direction modulations consistent with each monkey’s intact V1 initial direction biases (Fig. 8D-I), and consistent with the modeling results of Figs. 6, 7. This happened even though the saccade rate inhibition was abolished (Figs. 2, 3). For example, in both monkey F and the left V1 condition of monkey A, saccades after stimulus onset were weakly biased in the direction of the appearing stimulus. Naturally, there was no sharp oscillation in directions, but this is consistent with a lack of temporal resetting in the rate curve. That is, even though there was no temporal resetting, the impact of a latent visual signal appeared on saccades whenever these saccades occurred; it is the temporal resetting that sharpens the direction bias in time. For the right V1 condition of monkey A, there were again trends in the same direction as this monkey’s direction biases in the intact case; that is, there was still a similarity between intact and inactivation effects (but no statistical significance was reached in this case; there was also no sharp oscillation in the inactivation case). These observations validate the modeling results of Figs. 6, 7: rate inhibition can be lost while still seeing evidence of very weak direction biases (even without permanent lesions). For stimuli outside of the affected visual field, the saccade direction oscillations were all consistent with the intact situation (Fig. S13). Moreover, with saline injections, the direction oscillations in the injected hemifield were again similar to the normal situation (Fig. S10C, D, G, H). We obtained similar results with the white stimuli, as well as the memory-guided saccade task context.

Therefore, in all, even with muscimol inactivation, there was still evidence for a very weak latent visual signal in fixational saccade directions, but not saccade rates, consistent with predictions from the lesion and computational studies described above.

## Discussion

Exogenous sensory events inevitably and reflexively reset ongoing oculomotor activity (78), resulting in short-latency saccadic inhibition. Although the dynamics and functional implications of saccadic inhibition have been extensively studied, the underlying mechanisms mediating them have remained elusive (104). Popular models rely on lateral inhibition in oculomotor structures, such as the SC and FEF (85, 87, 89, 94, 100–103).

However, it has been shown that reversible inactivation of these structures alters some aspects of saccadic inhibition but does not eliminate the early inhibition phase. For example, muscimol SC inactivation affected microsaccade directions but not microsaccade rates (105). Similarly, FEF cooling influenced the late rebound phase after inhibition, but the early inhibition was unaffected (106). Ultimately, it was first proposed by Hafed and Ignashchenkova (98) that saccadic inhibition might result from short-latency sensory responses in the lower brainstem, which functionally raise the threshold for saccade generation and countermand saccades (78, 142).

However, regardless of the precise oculomotor mechanism, the question remains: which visual pathway dominates saccadic inhibition? In primates, the major candidate for such a role is the geniculostriate pathway, which brings visual information from the retina to the LGN and then V1. This pathway accounts for about 90% of the retinal ganglion cell projections exiting the eyeball (7), and our results are consistent with its dominance – we found that loss of V1 renders saccadic inhibition statistically unobservable. Similarly, based on human psychophysical experiments, it has been shown that saccadic inhibition requires sensory awareness (understood as an explicit perceptual detection) and is eliminated when orientation specific adaptation takes place (95, 96). These behavioral studies again give credence to our observations, although a direct role for V1 in sensory awareness is not necessarily as straightforward to directly infer from psychophysics alone (4), and the primate brain has other, alternative, visual routes that can bypass V1 and bring visual information to the lower brainstem and mediate saccadic inhibition.

The problem of inferring which pathway is most relevant for saccadic inhibition was historically rendered even more complex due to evidence from permanent V1 lesions (46, 47, 73, 74, 125, 143). With such lesions, both humans and monkeys can exhibit residual visual abilities that can help guide eye movements, resulting in a general consensus that either direct retinotectal projections or other projections bypassing V1 must be normally active in the intact brain. In fact, Rafal et al. (13) showed that in hemianopic patients not showing signs of blindsight for other tasks, visual distractors in the blind field could successfully inhibit saccades to targets in the intact field. These results, which are directly relevant to our experiments, were interpreted as suggesting the involvement of retinotectal pathways. Nonetheless, and especially given the almost-complete loss of SC activity seen with large-scale reversible LGN inactivation (144), the above-mentioned consensus needed serious revisiting. This is what we achieved with our experiments, and specifically by our comparisons between transient and permanent V1 inactivations. From these comparisons, we conclude that our observations are only partially consistent with the idea that retinotectal and other projections are normally strongly implicated in behavior in the intact brain. In fact, when measuring saccade rates, the dominance of V1 is decidedly unambiguous based on our transient inactivation experiments; these experiments were presumably transient enough to not allow long term brain plasticity to occur. At the same time, when measuring saccade directions, we did manage to find weak evidence for an influence of alternative visual pathways even in the transient inactivation experiments. This was also the case in our computational model, assuming an independence between rate and direction effects, and the lesion experiments. Thus, there is now a very pressing need to further investigate under what other sensory conditions can the alternative visual pathways bypassing V1 be more functionally impactful on behavior in the intact brain.

Our permanent lesion experiments were also instrumental in concluding which aspects of saccadic inhibition (e.g. saccade rate versus saccade direction modulations) might be mediated by alternative visual pathways in the intact brain. This is because these lesion experiments revealed that recovery of the inhibition phenomenon is indeed possible, and that saccade direction modulations are also observable. Of course, we had variable results across the three lesioned monkeys. This could be due to individual differences in the long-term brain changes that might happen after the lesions. For example, there could be differences in the rearrangement that takes place of LGN projections to the cortex (77).

Indeed, human hemianopia patients are widely variable and do not all develop blindsight (4, 57). In our case, the three monkeys were also variable in their behavior before the lesions, and the stimuli for some of them may not have been ideal to study saccadic inhibition (e.g. with gradual luminance changes; Methods). Nonetheless, their use in our study was a deciding factor in helping interpret our transient inactivation results.

Finally, it is worthwhile to view our results in light of other types of apparently reflexive ocular responses; doing so renders our observation of a dominance for the geniculostriate pathway in saccadic inhibition plausible, albeit still remarkable. For example, when a large visual field image suddenly moves coherently in one direction, smooth ocular following responses are initiated within ∼50 ms from motion onset in monkeys (145–147). Lesion experiments in the medial superior temporal area (MST) were shown to cause a large, and sometimes complete, reduction of even the shortest-latency component of these ocular following responses (148). This suggests that the brainstem circuits mediating ocular following responses depend on the cortex. On the other hand, deficits in area 17 of the cat were less complete (149). It would be interesting in the future to investigate whether V1 inactivation, like in our experiments, also eliminates the even smaller ocular drift responses recently described in monkeys (150, 151), which also have very short latencies from stimulus onset.

In all, our results recast some classic views on interpreting even the most reflexive of oculomotor behaviors, and they motivate further pathway investigations ultimately leading to eye muscle control.

## Acknowledgements

We were funded by the Deutsche Forschungsgemeinschaft (DFG; German Research Foundation) through the following projects: SPP 2411 Sensing LOOPS: Cortico-subcortical Interactions for Adaptive Sensing (project numbers: 520617944 and 520283985, HA 6749/11-1); BU4031/1-1; SPP2205 Evolutionary optimization of neuronal processing (project numbers: 430158665 and 430157666, HA6749/3-2); SFB 1233 Robust Vision (project number: 276693517); and BO5681/1-1. We were also supported by grants from the Japan Society for Promotion of Science (project numbers: 22H04992 and 22220006), from AMED Japan (project 24wm0625201h0001), as well as through internal Funding from WPI-ASHBi of Kyoto University (to author TI).

## Author contributions

TM, MPB, YY, TZ, ZMH performed the reversible inactivation experiments. TM, ZMH performed the data analyses, and the computational modeling. MY, TI, XY performed the lesion experiments. TM, ZMH wrote the first draft. All authors read and approved the final manuscript.

## Methods

### Experimental animals and ethical approvals

For the reversible inactivation experiments, we collected eye movement data and recorded, as well as inactivated, V1 neuronal activity from two adult, male rhesus macaque monkeys (F: 14 years, 13 kg; A: 12 years, 9.5 kg). Both monkeys were previously prepared for behavioral and neurophysiological experiements (109, 152–155). Briefly, they were implanted with scleral search coils for measuring eye movements using the magnetic induction technique (156, 157), and they were also implanted with recording chambers over the occipital lobe.

For the permanent lesion experiments, two adult monkeys’ data (specifically, from monkeys Ak and Tb) were obtained from a previous publication (63). Briefly, monkey Ak (Macaca fuscata) was male and weighed 9 kg; monkey Tb (Macaca fuscata) was female and weighed 6.5 kg. The third adult monkey (S; Macaca fuscata) was male and weighed 4 kg; the data from this monkey were obtained from (158).

The reversible inactivation experiments were approved by the regional governmental authorities of the city of Tübingen (Regierungspräsidium Tübingen, Germany). For monkey S, the experimental procedures were approved by the Committees for Animal Experiments at the Graduate School of Medicine in Kyoto University and at the National Institute of Natural Sciences (Japan). For monkeys Ak and Tb, the experiments were approved by the Committee for Animal Experiment at the National Institute of Natural Sciences (Japan).

### Laboratory setups

For the reversible inactivation experiments, the experimental setup was the same as previously reported (82, 159, 160). Briefly, the monkeys sat in a dark room facing a CRT display centered at their eye level. The display subtended ∼30 deg horizontally and ∼23 deg vertically and was placed 72 cm in front of the animals. All stimuli were presented against a gray uniform background of 26.11 cd/m^2^ luminance. Experimental control was achieved with a modified version of PLDAPS (161–163), interfacing with the Psychophysics Toolbox (164) and an OmniPlex data acquisition system (Plexon, Inc.). Injectrodes (linear multi-electrode array probes with a fluid channel) were obtained from Plexon, Inc., and they were positioned in the cortex using a NAN Microdrive (NAN Instruments, ltd). Muscimol injection (described in more detail below) was performed with the aid of a micro-injection pump (‘11’ Plus Single Syringe, Harvard Apparatus, Inc.).

The permanent lesion experiments on monkeys Ak and Tb were performed in the laboratory setup described in ref. (63).

Monkey S sat in a dark room with head fixed facing a monitor (Diamondcrysta WIDE RDT272WX [BK]; Mitsubishi) centered at eye level. The monitor was placed at 60 cm from the eyes of the monkey. All stimuli were presented against a dark uniform background of 9.28 cd/m^2^ luminance, and also using the Psychophysics Toolbox. Eye movements were recorded by a video-based eye tracker (EyeLink 1000 PLUS; SR Research) at a sampling rate of 500-1000 Hz.

### Experimental procedures

#### Behavioral tasks for the reversible inactivation experiments

*Receptive field (RF) mapping task*. We used RF mapping procedures like described previously (116, 165, 166). Briefly, in the beginning of each trial, the monkey foveated a small black spot in the center of the display (0.18 x 0.18 deg) and was required to maintain fixation throughout the trial. After a period of 700-900 ms of stable fixation, a white visual stimulus (0.2 x 0.2 deg) was presented. We manually chose the location of the stimulus on the upcoming trial based on the RF estimates derived from the electrode position. On each trial, we observed multi-unit activity (MUA) evoked by the stimulus on the attended channels, both visually using the OmniPlex data acquisition software and aurally via speakers. This allowed us to determine the stimulus locations eliciting the strongest MUA responses across the available channels. Typically, we collected a few hundred trials of this task per session.

Later, we verified these locations ofline and plotted the RF maps (see *Offine confirmation of the inactivated area*).

*Delayed saccade task*. To estimate the region of the induced cortical blindness, we used a delayed visually-guided saccade task. In each trial, the monkey fixated a small white fixation spot (0.18 x 0.18 deg) of 79.9 cd/m^2^ luminance. After a random period of 300-700 ms, a white visual target (0.18 x 0.18 deg) was presented and stayed on the display. The monkey was required to keep gaze at the fixation spot until it disappeared (the go signal) and then to foveate the target with a saccade. The interval between the target presentation and the go signal varied from 500 to 1000 ms and the grace period for target acquisition was 500 ms.

The trial was rewarded if the response saccade landed within 3 deg radius window. On each trial, the target location was automatically chosen from a grid of potential locations with resolution of 2 x 2 deg. However, on some trials, we manually placed the target in the predicted scotoma region and monitored the monkey’s behavior. In these trials, the monkey performed a guessed saccade, if any, to some random location in the visual field, so that the error between the target location and the saccade endpoint was substantial, and the monkey was not rewarded. Critically, we enforced a no-repeat strategy across all trials. That is, if the monkey failed to foveate the target, or to get the final reward of the trial for any other error (such as a premature fixation break) we never repeated the same trial again; a random position from the above-mentioned grid layout was chosen for the next trial. This way, we minimized the possibility that the monkey would learn the dimensions of the rewarded window and attempt to guess target location if it was not perceived due to the scotoma. As a result, the error in the saccade endpoint position was pronounced enough to allow us to estimate the scotoma area by eye. We performed this estimation on the go and placed the visual stimulus in the subsequent tasks accordingly. The scotoma area was later verified ofline with a saccade error map analysis (see *Offine confirmation of the inactivated area*). We analyzed 118-317 trials of these tasks per session in monkey F (3439 trials in total) and 113-375 trials per session in monkey A (4132 trials in total).

*Fixation task*. The monkey always had to fixate on a black fixation spot of 0.18 x 0.18 deg dimensions in the center of the display. After a random period of fixation (300-500 ms), a visual stimulus of 0.36 deg diameter was presented in a predefined location (as per the above) in the scotoma region. In other trials, the same stimulus appeared in the opposite (intact) visual hemifield, by flipping the sign of its horizontal position relative to the scotoma location. In some sessions, we also had control trials (3.2% of the trials in the session), in which no stimulus was presented at all (we experimentally marked a fictive onset using the same timings as with the real stimulus onset). In all trials, the monkey had to keep fixating until trial end, which occurred 500-800 ms after the offset of the visual stimulus (which was itself on for 100 ms). We manipulated the luminance polarity of the stimulus and its Weber contrast, so that the trials were equally divided across the following conditions: 100% Weber contrast black, 10% Weber contrast black, and 100% Weber contrast white. We also collected intact V1 sessions, which were run on other days than the inactivation sessions days. With this task, we collected 34 sessions in monkey F (17 inactivation sessions and 17 matching intact sessions) and 20 sessions in monkey A in the right V1 (10 inactivation sessions and 10 matching intact sessions). We also collected 26 sessions in monkey A’s left V1 (13 inactivation sessions and 13 matching intact sessions). Additionally, we collected 37 extra intact sessions in this monkey. In monkey F, we analyzed 403-1697 trials of the task per inactivation session (10361 analyzed trials in total) and 435-1708 trials per intact session (10116 analyzed trials in total). In monkey A, the trial numbers were 900-2001 trials of the task per inactivation session and 171-1743 trials per intact session. In monkey A, we analyzed 8294 inactivation trials and 22615 intact trials in right V1 and 5696 inactivation trials and 4044 intact trials in left V1.

*Memory-guided saccade task*. In the beginning of the trial, the monkey was required to fixate on a white fixation spot (0.18 x 0.18 deg) in the center of the display. After a random period of fixation (300-700 ms), a visual stimulus (0.36 deg diameter; 100% Weber contrast black) was presented in the predefined location in the scotoma region (∼18% of trials) or in a symmetric location in the opposite (intact) visual hemifield (∼18% of trials). The stimulus stayed on the screen for 50 ms, and the monkey was supposed to remember its location while keeping fixation at the center. After a random period of 500-1000 ms, the fixation spot disappeared, and the monkey had to perform a saccade towards the remembered location. After the monkey entered the saccade acceptance window, the target reappeared, and a reward was provided a few hundred milliseconds later. We set the saccade acceptance window to 3 or 4 deg radius around the target location to make it easier to execute a saccade towards the required location even if the monkey could not see the target; even with an intact V1, memory-guided saccades are also less accurate than visually-guided ones (Fig. S7). We trained the monkeys to perform the task in a way that they learned that if no target was visible to them, the saccade should be performed to the portion of the screen representing the scotoma region (one of the two lower quadrants). We also had control trials (∼9%), in which no stimulus was presented at all, and the monkeys were still required to saccade to the same scotoma area (or a corresponding region in intact sessions). As above, the intact sessions were run on separate days. We collected 14 sessions in monkey F (7 inactivation sessions and 7 matching intact sessions) and 9 sessions in monkey A in right V1 (5 inactivation sessions and 4 matching intact sessions; the reason it is 4 is that two inactivation sessions had the same target location, so we simply collected more trials on one of the intact sessions to account for that). In total, we analyzed 1642 inactivation trials and 1605 intact trials in monkey F; in monkey A, these numbers were 1961 and 1520 trials, respectively. In all cases, for the purposes of this study, we were primarily interested in studying the fixational saccades occurring around initial stimulus presentation and before the go signal for the final memory-guided saccade.

#### Behavioral tasks for the permanent lesion experiments, monkeys Ak and Tb

For the permanent lesion experiments (monkeys Ak and Tb), data were collected under four conditions: pre-lesion (3–4 weeks prior to the surgery) and post-lesion (4–19 weeks post-surgery) for each animal. The trial structure matched that of the memory-guided saccade of monkeys A and F, except that most trials utilized a gradual target onset (contrast ramp of 40– 240 ms duration). In the pre-lesion sessions, stimuli appeared at 4 or 6 radial locations, at an eccentricity of 10 deg. For monkey Tb, the visual targets had either an immediate onset (∼32%) or a 90–180 ms contrast ramp, with a 1.5 deg diameter and brightness of 2.8–27 cd/m^2^. For monkey Ak, all trials involved a target with 180 or 240 ms contrast ramps, and with 0.45 or 0.90 deg diameter. In the post-lesion sessions, the parameters of the target were further refined per the requirements of the original experiments. For monkey Tb, we used a fixed 180 ms contrast ramp and 1.5 deg target diameter. For monkey Ak, we used 40 or 240 ms contrast ramps with brightness levels of 0.55–27 cd/m^2^ (0.45 deg diameter in 13% of the trials, and 1.5 deg diameter in the remainder). In all experiments, the background luminance was 0.18 cd/m^2^. In some sessions, targets were restricted to either the affected or intact hemifield to focus the data collection strategy. Analysis included only rewarded trials, and we analyzed all fixational saccades occurring before fixation spot offset in the memory-guided saccade task. For monkey Tb, we analyzed 377 pre-lesion and 6503 post-lesion trials; for monkey Ak, these numbers were 3952 and 2592 trials, respectively.

#### Behavioral tasks for the permanent lesion experiments, monkey S

The task was a 4-alternative forced-choice visually-guided saccade task. Each trial began with a 300-1000 ms maintenance of central fixation, followed by presentation of four peripheral targets. After a 500 ms pre-go delay, a go cue instructed the monkey to make a saccadic choice. Following correct choices, confidence was expressed by whether the monkey maintained or voluntarily broke central fixation during a post-choice fixation period, which was linked to the reward magnitude. Again, here our purpose was to analyze fixational saccades before the go signal and after eccentric stimulus appearance. Only rewarded trials (3202) were included in the analysis.

After the V1 lesion, the monkey was trained on the same task. The visual parameters of the display were as follows. The central fixation spot had a diameter of 0.6 deg and a Michelson contrast of 0.5. Target directions from the horizontal were at 126, 162, 198, and 234 deg in the intact visual field, and 306, 342, 18, and 54 in the affected visual field. The targets were all at an eccentricity of 10 deg, and all had a diameter of 0.3 deg. The targets had 0.0001 to 0.8 Michelson contrast in the intact visual field, and 0.01 to 0.95 Michelson contrast in the affected visual field.

The behavioral data in the present study were obtained 16-20 months after V1 lesioning. We also confirmed that the affected visual field had been already largely recovered at the time of data collection, since the hit rate reached over 90% with target contrasts (Michelson) higher than 0.6; and over 65% with target contrasts higher than 0.2.

#### Muscimol injections

In all reversible inactivation sessions, we administered injections of the GABA-A agonist muscimol using linear multi-electrode arrays with a fluid channel (FC) from Plexon. We collected 34 sessions with 16-channel V-Probes (7 sessions: 100 μm spacing; FC between channels 7 and 8; 27 sessions: 150 μm spacing, FC between channels 8 and 9) and 6 sessions with 8-channel S-Probes (200 μm spacing, FC between channels 8 and 9). The same multi-electrode arrays were used to determine V1 RF locations prior to the injections (and thus to predict the expected region of deficit), and also to confirm tissue silencing upon muscimol delivery (Fig. 1B). We varied the inactivation site across sessions, but it was always restricted to the dorsal part of V1 representing the lower visual field (Fig. 1C; Table S2).

To avoid potential distortion or silencing of neuronal activity due to muscimol being at the interface of the fluid channel with the tissue, we estimated the pre-injection RF’s immediately after penetrating V1 but before going deeper into the tissue; that is, we monitored intact V1 state before injection using the deepest one or two contacts of the electrode array. This estimation was performed online using MUA from the active channels, while simultaneously running the RF mapping task. We later confirmed the RF’s online (see below).

We performed the injections with a Hamilton syringe (1800 series, gastight, 10 μL) mounted on the micro-injection pump and connected to the probe via transparent polythene tubing (0.28 μm inner diameter). The Hamilton syringe and the adjacent portion of the tubing were filled with saline (0.9% NaCl solution) mixed with food colorant serving as a visual marker and separated by an air bubble from the muscimol. The muscimol was dissolved in saline (concentration: 10 mg/mL) and filled the remainder of the tubing and the probe. We used the displacement of the air bubble along the tubing to confirm the volume of the muscimol delivered per injection.

We infused muscimol at multiple depths in V1 by first advancing the probe to deeper V1 layers and then retracting it to more superficial layers. Each injection pulse lasted for 1-2 min, and muscimol was delivered with a rate of 0.1-0.2 μL/min in each pulse. The waiting time between injection pulses was 2 min, during which some passive leakage of muscimol could sometimes be observed. The overall injection period for a session lasted 20-120 min, and the total volume of muscimol administered was 1.13-2.78 μL (see Table S2). We then waited for at least an additional 20 min to confirm that neuronal activity was silenced. The minimum duration from the injection start to the start of the data recordings was 53 min. The probe remained in the brain until the end of the session to monitor the neuronal activity, but we sometimes also retracted it after the injection. Overall, we conducted 17 inactivation sessions in monkey F (right V1) and 23 sessions in monkey A (10 and 13 sessions in the right and left V1, respectively; Table S2).

In each monkey, we also ran a sham inactivation session, in which we administered saline instead of muscimol. In these sham sessions, we followed the exact injection protocol as described above, with the only exception being that the probe and the adjacent tubing were filled with pure saline solution. The amount of saline infused was 1.28 μL in monkey F and 1.19 μL in monkey A.

#### Lesion procedures

For monkeys Ak and Tb, we used data from prior publications describing the lesion procedures (63, 167).

For monkey S, we used data from a prior publication describing the lesion procedures (158). Briefly, the left V1 was surgically removed by aspiration under anesthesia (details of the surgery have been described elsewhere (62, 167)).

### Data analysis

We detected saccades and microsaccades using our standard procedures (147, 168). Only rewarded trials were analyzed, except for the delayed saccade task, in which we were interested in the saccade endpoint errors (see below). All analyses were performed in MATLAB 2020b and 2025a (The MathWorks, Inc, USA) using custom scripts. For each condition, we pooled the data across all sessions; we did this because we have evidence that saccadic inhibition does not adapt to repeated trial exposures (79, 82). The monkeys’ data were analyzed separately; also, the left and right visual fields were treated independently in monkey A’s case. A fraction of the current data (namely, the black 100% contrast condition in the intact V1 fixation task) was analyzed for a recent behavioral report (82), but no inactivation or lesion data were previously analyzed.

#### Offine confirmation of the inactivated area

To confirm the V1 RF’s, we extracted MUA from active channels. Specifically, we band-pass filtered the wideband signal (4^th^-order Butterworth, 750-5000 Hz), rectified it, and then low-pass filtered it (4^th^-order Butterworth, 500 Hz) to obtain the signal envelope. The resulting signal was downsampled (from 40 KHz) to 1 KHz. We then averaged pre-stimulus activity across trials in the interval from-100 to-1 ms relative to stimulus onset and subtracted this value from the MUA signal of the whole trial. Next, for each trial, we calculated the average visual response elicited by the stimulus in the interval from 20 to 110 ms after stimulus onset and converted it to a z-score by dividing it by the baseline standard deviation. After this, we plotted the z-scored visual response strength as a function of stimulus location. For that, we divided the visual field into a 2-D grid with resolution of 0.05 x 0.05 deg and fitted a continuous surface using the natural neighbor interpolation method. Finally, we applied a z-score threshold of 1.65 (corresponding to 95% one-tailed confidence intervals) and set the interpolated surface values below it to zero. This procedure allowed us to confirm that pre-injection RF locations were consistent with our planned V1 site entry, and also consistent with our subsequent evaluation of the scotoma.

To confirm the area of inactivation, we plotted the saccade error maps derived from the delayed saccade task. In this task, we included not only the rewarded trials but also the trials in which the disappearance of the fixation spot (the go signal) prompted the monkey to make a saccade that landed outside of the rewarded target window. In any given trial, we calculated the Euclidian distance between the eye position at the response saccade endpoint and the actual target location. We then plotted the error magnitude as a function of actual target location. For that, we binned the target locations into 0.25 x 0.25 deg grids, interpolated the missing values using the natural neighbor method, and visualized the obtained continuous surface (e.g. Fig. 1G-I). We also obtained an estimated contour surrounding the induced scotoma (e.g. Fig. S1). To do this, we plotted only those areas where the error exceeded 1.5 deg. One inactivation session in monkey F was not analyzed ofline due to a technical error during the file saving process; however, we were still able to locate the scotoma region by online saccade error monitoring while recording the task, which was also consistent with the online estimation of the V1 RF locations.

#### Microsaccade rate analyses

For tasks with response saccades (e.g. memory-guided saccade task and the lesion monkey behavioral tasks), we only analyzed fixational saccades before the go signal for the response saccade (response saccades with muscimol were analyzed in Fig. S7 and with permanent lesions in other publications). We estimated microsaccade rate using an approach similar to those we used earlier (80–82). Specifically, on any given trial, we calculated the number of microsaccades that occurred within a 50 ms time window; we moved this window by 2 ms and repeated the procedure to obtain the full time-course. We then divided the obtained numbers by 50 ms to obtain a rate of microsaccades per second. Finally, for any given condition, we aligned the trials relative to the time of the stimulus onset, averaged the microsaccade rate for each time point across trials, and estimated the pointwise standard error of the mean across trials.

To statistically assess the effects of experimental manipulations on the microsaccade rate signature in the fixation and memory-guided saccade tasks, we first used cluster-based non-parametric permutation tests (80, 169–171) and tested each condition of interest against the control (no-stimulus) condition. To increase the number of trials in the control condition, we pooled data from the intact and inactivation trials, since the trial structure was identical and no stimulus was presented in either case. Prior to this, we ensured that there were no significant differences between the control conditions in the two datasets (Fig. S2D, E). We also merged the left and right V1 intact sessions in monkey A into a single dataset (while still accounting for stimulus hemifield in individual condition analyses), since these trials were identical (that is, in the intact trials, whether the right or left visual stimulus was in the scotoma during reversible inactivation did not change the fact that the intact sessions had both a right and left visual field stimulus presentation). Because we aimed to maximize sensitivity of our tests to potentially subtle effects – as might be expected in the absence of V1 input – we restricted our analyses to the time windows in which microsaccadic inhibition and rebound were known to occur in similar tasks in our animal model (79–82, 98, 104, 105), and we ran separate one-sided tests on each of these windows. For the microsaccadic inhibition interval, the time window was set to 30 to 220 ms relative to the stimulus onset (or to a corresponding event marker in the control condition); for the post-inhibition rebound interval, it was set to 150 to 300 ms.

For each comparison, we computed a pointwise test statistic across the designated time interval, defined as the difference between the mean instantaneous microsaccade rate in the condition of interest and that in the control condition. We then pooled all trials from the two conditions into a single set, randomly reassigned their labels while maintaining the original trial ratio, and recomputed the test statistic. We repeated this permutation procedure 10000 times and stored the resulting differences as the null distribution. Next, when we ran the test on the inhibition epoch, we identified all continuous time periods in which the observed data fell below predefined significance level (alpha = 0.05), i.e. below the lower 5^th^ percentile of the null distribution. For the rebound epoch, we identified the continuous periods in which the observed data exceeded the 95^th^ percentile of the null distribution. Those periods were thus defined as the significant clusters in the real data.

To account for multiple comparisons, we applied the same clustering procedure to each permutation sample and recorded, for each permutation, the size of the largest cluster. We chose the cluster size over the cluster mass (i.e. the sum of the magnitudes of all test statistics within a cluster) as the cluster-level statistic because it was sensitive to weak but temporally stable effects, which might otherwise be masked by transient, high-amplitude accidental fluctuations in the microsaccade rate. Finally, for each cluster in the observed data, we calculated its Monte Carlo p-value (*p_MC_*) as the proportion of permutation clusters whose cluster-level statistic was equal to or greater than the cluster-level statistic of the experimentally observed cluster. Clusters with *p_MC_* < 0.05 were considered significant.

When we did not have a control condition, as in the permanent V1 lesion datasets, we ran one-sample cluster-based permutation tests, again separately for the inhibition and rebound epochs. For any given condition, we computed a point-wise difference between the observed data in the predefined time window and the average pre-stimulus microsaccade rate, which was calculated across trials on the interval-50 to-1 ms relative to the stimulus onset. In this case, the null distribution was obtained by randomly flipping the signs of the trials (i.e. by multiplying each trial by either 1 or-1) and repeating this procedure for 10000 times. The remainder of the analysis, such as identifying the significant clusters and correcting for multiple comparisons, was performed in the same way as for the two-sample permutation tests described above.

To directly compare two conditions with rate modulations (e.g. intact versus inactivated microsaccade rate signature), we ran again the two-sample cluster-based permutation tests. Since we had no informed predictions about the timing or directions of effects to observe in this case, we applied the tests on the whole post-stimulus interval of 30 to 300 ms and considered the observed clusters to be significant if their test statistic was below the 2.5^th^ percentile (for negative clusters) or above the 97.5^th^ percentile (for positive clusters) of the null distribution. The remaining procedure was the same as described for the two-sample permutation tests above, with the exception that the critical alpha level for the Monte Carlo p-values was now set to 0.025. We applied this analysis when comparing the intact and inactivated V1 datasets against each other, and when comparing the intact trials with the saline test trials. Note that we did not directly compare the pre-lesion and post-lesion data with each other in monkeys Tb and Ak due to fundamental differences in the experimental procedures, such as the range of the stimulus contrasts used and the duration of the gradual stimulus appearance. In case of monkey S, we had no access to the pre-lesion data.

#### Microsaccade direction analyses

We used the same approach to estimate the microsaccade direction modulations as we introduced recently (82). Specifically, we computed the baseline-subtracted rate difference between microsaccades towards one or the other hemifield; this metric allowed us to account for any default idiosyncratic biases in microsaccade directions even before the stimulus was presented. To do so, we first classified microsaccades as being directed either towards or away from the stimulus location (hemifield-based), and we calculated the microsaccade rate separately for each of these classes, following the same rate calculation procedure described above. Next, we estimated the mean baseline rate for each class (in the interval from-50 to 0 ms relative to stimulus onset) and subtracted it, at every time sample, from its rate curve. Lastly, we obtained the rate difference curve by computing the pointwise difference between the ‘towards’ and ‘away’ curves. This metric being above zero indicated a bias towards the stimulus hemifield, whereas when it was below zero, it reflected a bias away from the stimulus hemifield.

To statistically access the effects of our experimental manipulations on microsaccade directions, we ran one-sample cluster-based permutation tests similar to ones we used for the rate statistics above. That is, we calculated the difference between the post-stimulus microsaccade rate and the mean pre-stimulus rate (-50 to-1 ms relative to stimulus onset), randomly flipped the trial signs 10000 times to generate the null distribution, identified significant clusters, and applied a multiple-comparisons correction. We looked for positive and negative clusters in the whole post-stimulus interval from 10 to 300 ms. To prevent potential masking by stronger biases in one of the directions, the tests for positive and negative clusters were performed separately, each treated as a one-tailed test with alpha level of 0.025.

When comparing directional modulations between two conditions (e.g. Figs. 8D-F, S10C, D, G, H, S13), we performed two-sample cluster-based permutation tests over the full post-stimulus interval (10 to 300 ms). The procedure was identical to that used to compare microsaccade rates across conditions other than the no-stimulus one. This analysis was applied to both the intact versus inactivated V1 datasets and the intact versus saline test datasets.

### Computational model

We implemented an even simpler version of our previous saccadic inhibition models (98, 138), which were generally framed after (172). The purpose here was to minimize parameter spaces to one single variable associated with reversible V1 inactivation.

The model simulated microsaccade generation as a repetitive rise-to-threshold process. In each simulation time step (1 ms), there was an accumulator process, *M*, the value of which could be updated. In the absence of any exogenous stimuli, *M* could be in one of three states: zero (numerically simulated as any value less than 1), rising towards the movement “triggering” threshold of 1000 with some rate of change *r_B_*, or decreasing back towards zero upon microsaccade “triggering”. Each simulated trial was 5000 ms long, and *M* started at a value of zero at the beginning of each trial. A rate of rise, *r_B_*, was picked at random for the current ensuing movement plan, from a normal distribution with mean 8 and standard deviation 2 (if the generated random number was negative or zero, we pegged *r_B_* for the current movement plan at 1). Thus, the dynamics of *M* were dictated by:

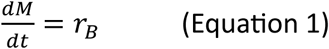

Once *M* crossed a threshold value of 1000, it underwent a rapid, exponential decay back to zero according to the following dynamics:

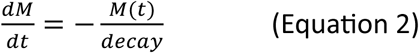

where *decay* was set to 7 ms. A saccade onset time was defined as the time of the threshold crossing plus an efferent delay of 20 ms. When *M* dropped to a value below 1, the process for the next microsaccade was to be started again. This entailed a short delay afferent processing delay of the next microsaccade target, randomly picked from a normal distribution with mean 95 ms and standard deviation 40 ms, after which Equation 1 proceeded with a new value of *r_B_*, again picked from the same normal distribution mentioned above. In both cases, if the random number generator returned a negative or zero value, we pegged the value of the respective parameter at 1.

To simulate microsaccade directions, the first movement of the trial was picked to have a random direction. Then, at the start of every subsequent movement plan (i.e. when *M* was supposed to rise again according to Equation 1), the current movement’s direction was picked at random from a normal distribution with a mean value of 180 deg plus the previous movement’s direction, and a standard deviation of 70 deg. This ensured that there was a general counter-phase relationship between successive microsaccades, as experimentally observed (98, 138).

If an exogenous stimulus appeared, it was supposed to countermand the currently ensuing microsaccade program. The countermanding process started after a sensory processing afferent delay of the exogenous stimulus, picked from a normal distribution with mean 30 ms and standard deviation 7 ms (once again, if the random number generator returned a negative or zero value, the value was pegged to 1). If *M* was rising after this processing delay, then the countermanding process immediately started acting on *M* by making *r_B_* a time-varying process:

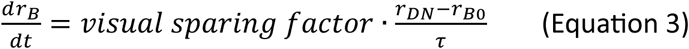

where *r_DN_* was set to-2.56, *r_B0_* was the initial rate of rise of *M* for the current microsaccade plan, and τ was 50 ms. If *r_B_* reached a value more negative than *r_DN_* before *M* either reached the movement triggering threshold (escape movement; failed countermanding) or <1 (successful countermanding), Equation 3 was stopped and *r_B_* remained at the value of *r_DN_* until *M* reached the quiescent state of <1 once again. The parameter *visual sparing factor* was our way to model the loss of visual drive by reversible V1 inactivation. When this parameter was 1, the model was intact; when it was less than 1, the countermanding efficacy was reduced.

If the exogenous stimulus was processed (i.e. after the sensory processing afferent delay mentioned above) and *M* reached the microsaccade triggering threshold nonetheless, then only for this microsaccade plan, the microsaccade direction was picked as having a mean value of 0 (assuming the exogenous stimulus was always defined as being in direction 0) and same standard deviation as above (70 deg). This stimulus-directed biasing by the exogenous stimulus reflected the influence of visual bursts in biasing the microsaccadic readout process in structures like the SC (141). Critically, it only happened for a single movement at the time of putative “visual bursts”. Moreover, the same *visual sparing factor* parameter was implemented here, such that its value reduced the likelihood that this microsaccade biasing was to happen by the stimulus. Specifically, for each such case of a stimulus-locked movement scenario, we sampled a random number from a uniform distribution between 0 and 1. If this number was greater than 1 minus *visual sparing factor*, the microsaccade direction was picked as above with a mean direction of 0 deg; if not, the microsaccade direction was just picked from the distribution with mean 180 deg from the previous microsaccade, just like all other microsaccades. Thus, a single parameter, *visual sparing factor*, influenced both microsaccade rate (Equation 3) and direction in the same way.

In each simulation, we ran 2000 model trials, with the exogenous stimulus onset occurring at a random time between 2000 and 3000 ms from trial onset. From these 2000 model trials, we calculated microsaccade rate on each trial. We did so by having a moving window of 50 ms, moved in steps of 2 ms. In each bin, we counted how many microsaccades occurred in the bin, and divided the number by the duration of the bin (50 ms). This gave us a rate estimate per trial. We then averaged all rate estimates across all 2000 trials. For directions, we again had 50 ms bins, stepped in steps of 1 ms relative to exogenous stimulus onset. In each bin, we calculated the proportion of microsaccades occurring in such a bin that had a direction towards the exogenous stimulus. This was defined in a hemifield manner; that is, if the microsaccade direction was towards the hemifield of direction 0, it was considered to be towards the stimulus, and if it had a direction in the other hemifield, it was considered as opposite. For Fig. 7G, H, we ran the full model (2000 trials each) for 25 times, each time for one particular value of the parameter *visual sparing factor*. Then, we quantified saccadic inhibition and direction biases by measuring microsaccade rate (Fig. 7G) or microsaccade direction biasing (Fig. 7H) in the interval 60 to 160 ms after stimulus onset. For Fig. 6C-F, we also fixed all parameters for the first microsaccade plan of the trial (i.e. not running the full stochastic model), just so that we could illustrate the effects of different exogenous stimulus onset times on the countermanding process for the exact same movement plan.

## Supplementary figures

**Figure S1.**
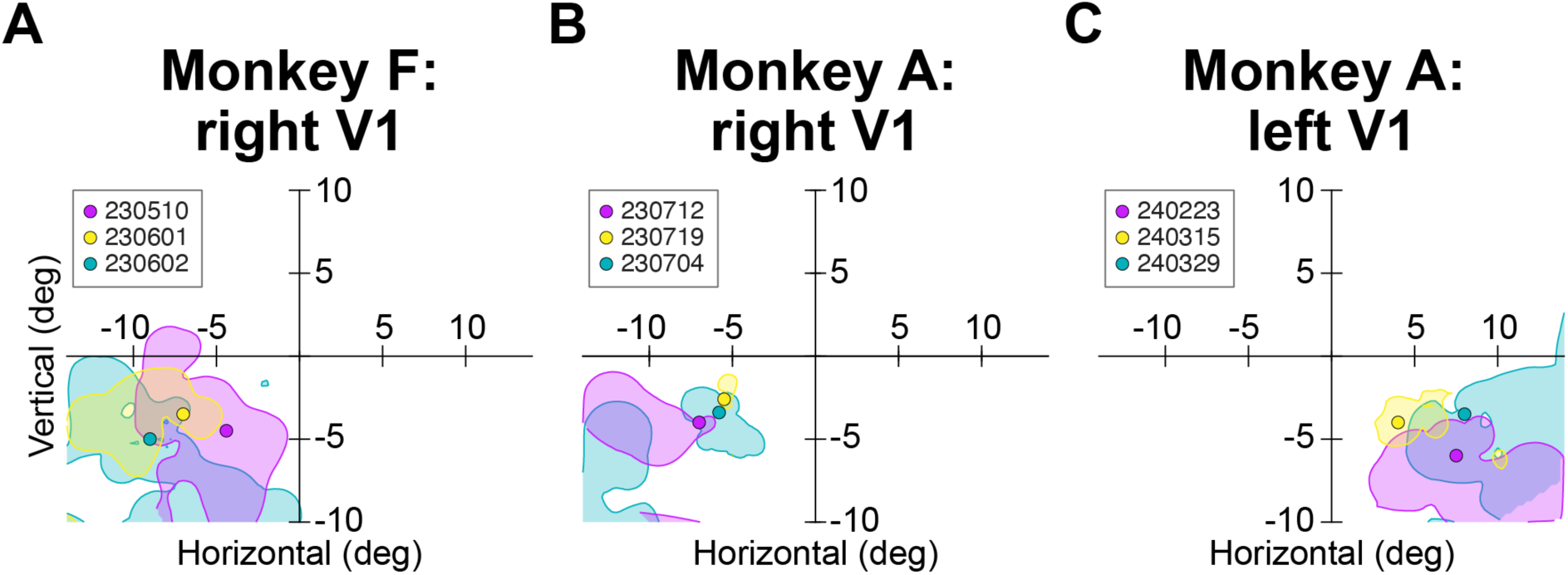
Additional example scotomas, and chosen visual stimulus locations within them, from each animal and each tested hemisphere. Each shaded region shows the locations of visual stimuli for which saccade landing errors after V1 inactivation (as in Fig. 1G-I) exceeded 1.5 deg. For each such region, the circle of the corresponding color indicates the location at which we placed the visual stimulus when we wanted it to appear within the cortically-blind region of the visual field. The numbers in the legends are session identifiers. In all cases, the stimulus was placed within the scotoma, and the scotoma was much larger than a single cortical column, confirming that we also inactivated all cortical layers (Fig. 1B and Methods). Note that the size of the circles shown here is not corresponding to the actual size of the stimulus that we used (Methods).

**Figure S2.**
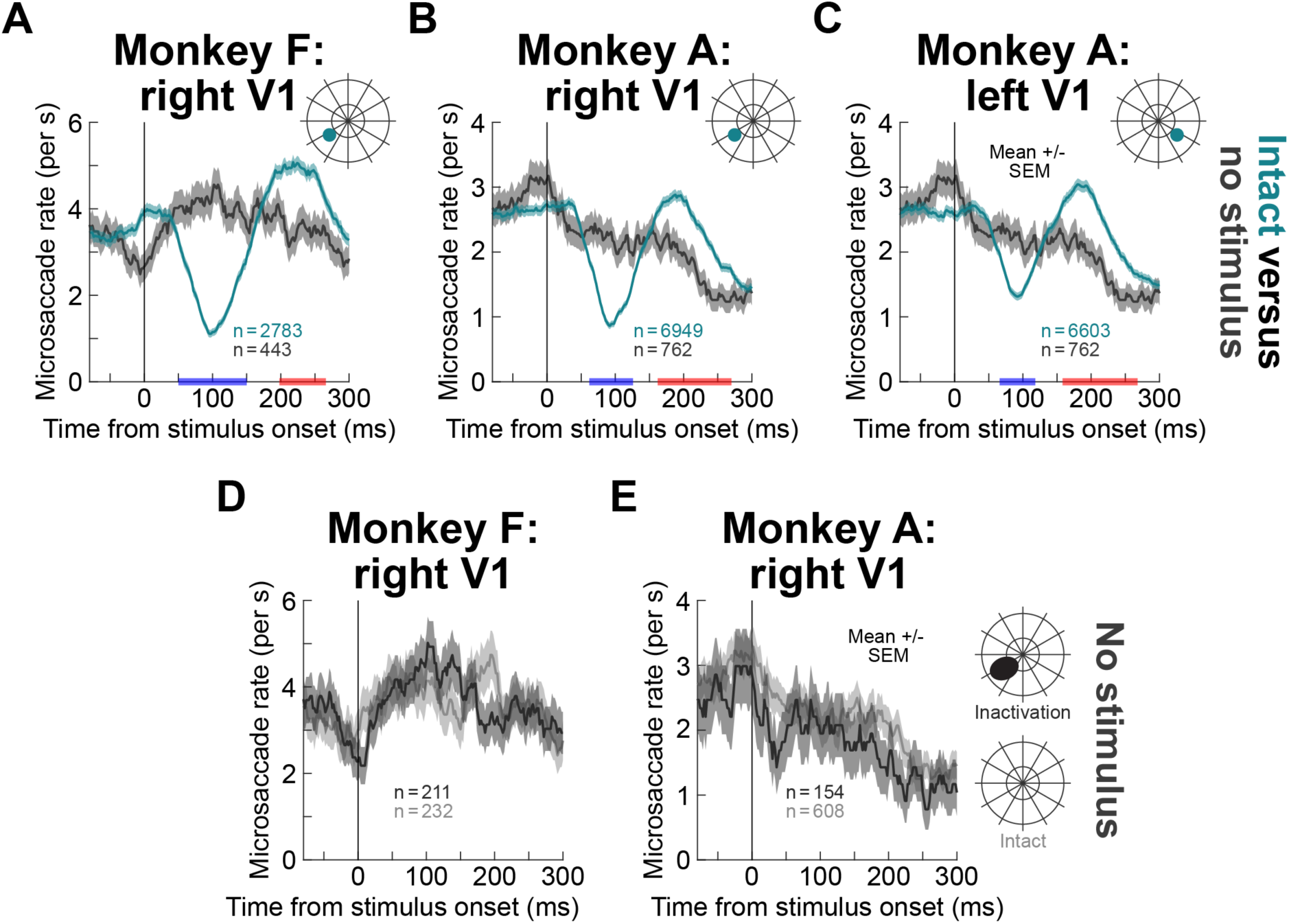
Intact V1 saccade rates are different from those with no visual stimulus appearing, and no-stimulus saccade rates do not depend on V1 inactivation state. (A-C) These panels are like Fig. 3A-C except that we now compared intact V1 data with a visual onset (greenish curves) to no-stimulus data (gray curves). Like with the inactivated V1 data of Fig. 3A-C, the rate curves differed within two distinct temporal epochs: saccade rates with a visual stimulus onset in intact V1 regions were lower than no-stimulus saccade rates at short latencies from stimulus onset (indicating saccadic inhibition; blue epochs on the x-axes); saccade rates with a visual stimulus onset in intact V1 regions were higher than no-stimulus saccade rates at longer delays (indicating rebound from inhibition; red epochs on the x-axes). Note how monkey A showed time-varying saccade rates even on no-stimulus trials. The results of the statistical comparisons giving rise to the red and blue epochs on the x-axes are included in Table S1. **(D, E)** There was no difference between no-stimulus saccade rates with and without V1 inactivation. For each monkey, we plotted saccade rate as a function of time from fictive stimulus onset in the no-stimulus control condition. There was no difference in such rate between having an intact or inactivated V1 region. In monkey A, the same time-varying decrease in saccade rate, in anticipation of trial end, was observed independent of the inactivation condition. Thus, inactivation did not alter the patterns of saccade generation of the monkeys in the absence of any visual stimulus onset. As a result, in **A**-**C**, as well as all other figures with no-stimulus data, we combined the no-stimulus trials of the intact and inactivation sessions for simplicity (as well as greater statistical robustness). Error bars denote SEM across trials.

**Figure S3.**
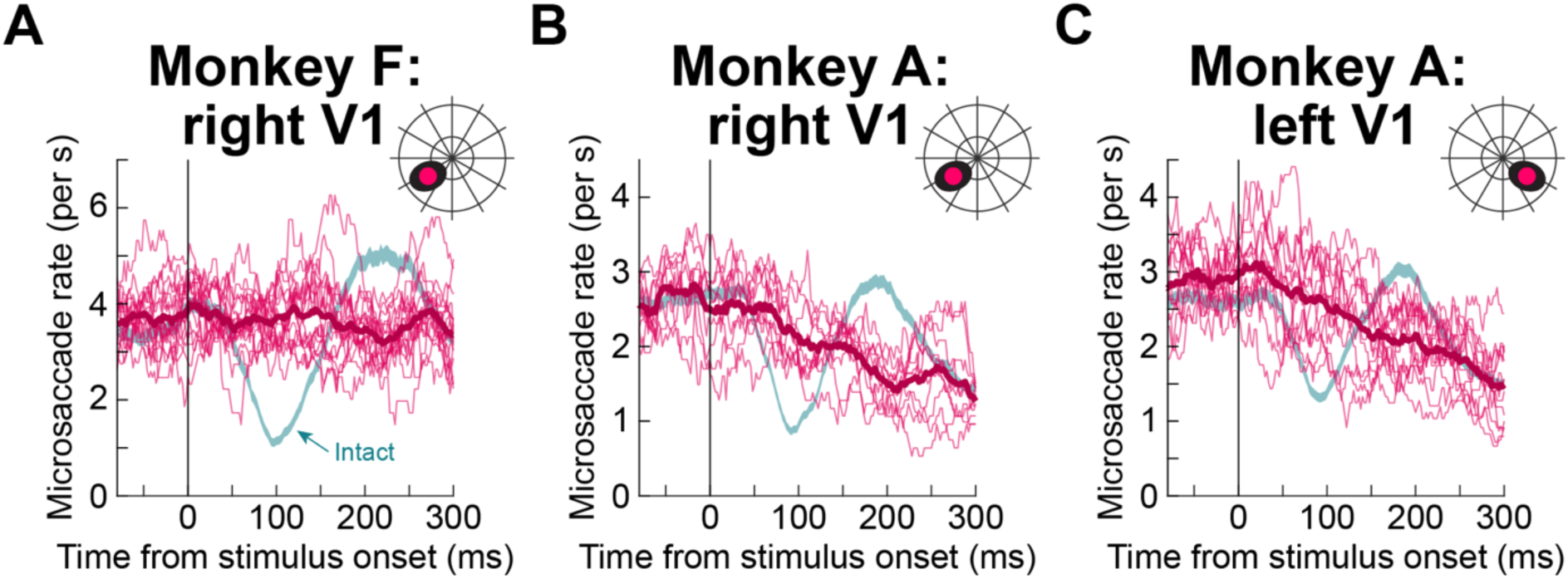
Consistency of the results of. **Fig. 3A-C across individual inactivation sessions.** This figure shows the inactivation data of Fig. 3A-C (reddish curves) but after separating these data across individual sessions. Each thin curve shows the average saccade rate from one inactivation session, and the thick reddish curve is the average across sessions. The greenish band shows the +/-SEM limits of the intact V1 data from Fig. 3A-C. As can be seen, in no single inactivation session with a visual stimulus appearing within the cortical scotoma was there evidence of saccadic inhibition even approaching the inhibition observed in the intact data (example single-session intact inhibition profiles can be seen in Fig. S10, and they are very different from those observed here). Thus, the results of Fig. 3A-C were highly robust across individual sessions. Note also that we have recently shown that saccadic inhibition does not adapt across trial repetitions, justifying the pooling of trials and sessions in our analyses (82) (also see (79–81, 83)). The total number of sessions is the same as in Fig. 1C.

**Figure S4.**
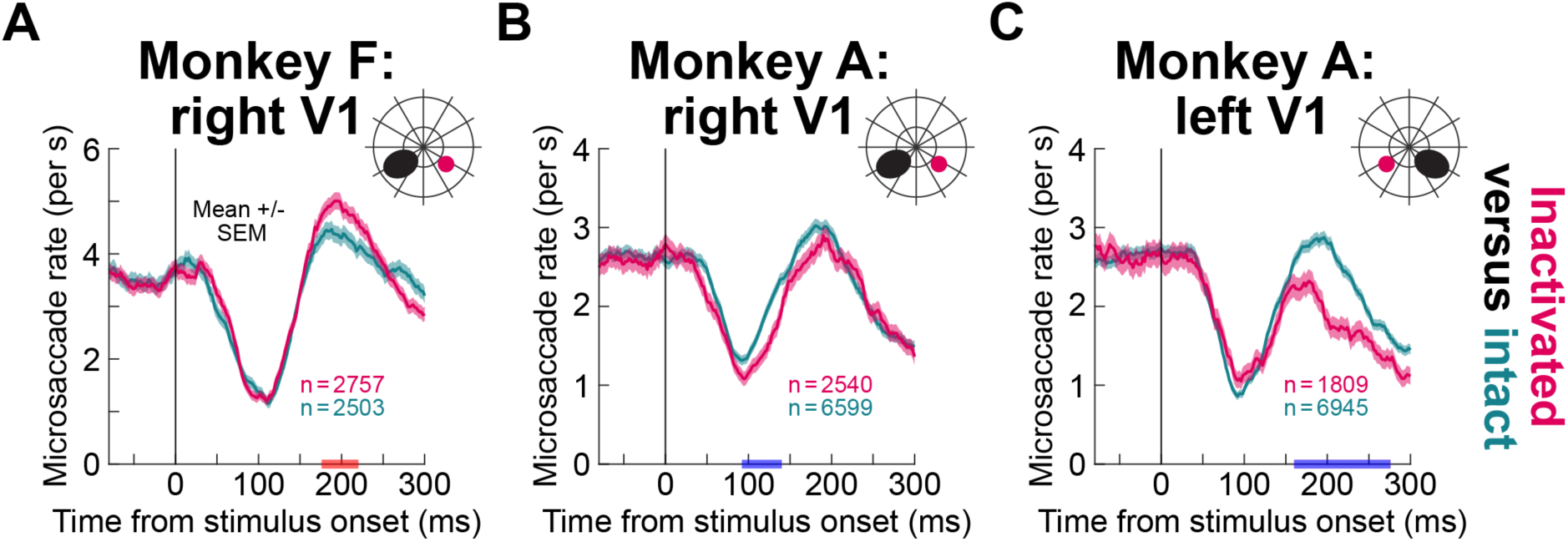
No impairment of saccadic inhibition during V1 inactivation with visual stimuli presented in the unaffected hemifield. (A-C) In each experiment, we also tested visual onsets in the hemifield opposite the scotoma location (Methods). Here, we plotted saccade rates from these trials under two conditions: greenish shows the rates without V1 inactivation, and reddish shows the rates with V1 inactivation (schematics above the data indicate the relative locations between the stimuli and the scotomas). As can be seen, the early inhibition phase (<100 ms from stimulus onset) was the same whether V1 was inactivated or not, suggesting that the results of Figs. 2, 3 were specific to stimuli being within the scotoma region. At later times after stimulus onset, there were significant differences that emerged between the rate curves in the two conditions in each monkey (indicated by colored epochs on the x-axes; Table S1). These differences could be expected based on the fact that post-inhibition rate rebound is a cortically-dependent phenomenon (80, 104, 106, 110–112); and, in monkey A, there were some visual field asymmetries in microsaccadic inhibition even without any cortical inactivation (82). Error bars denote SEM.

**Figure S5.**
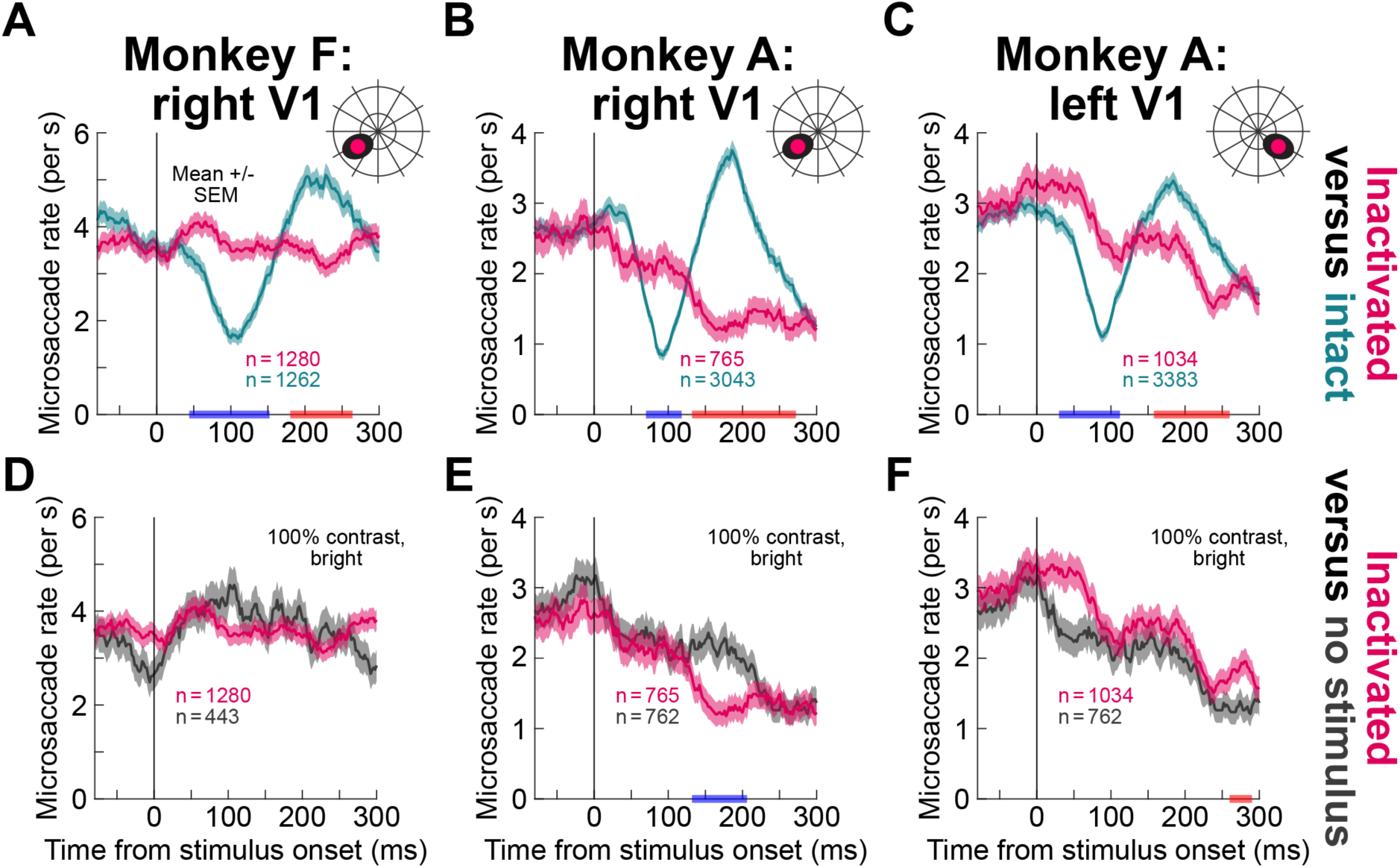
Similar results to. **Fig. 3, but with bright visual stimuli. (A-C)** Similar to Fig. 3A-C, but with 100% contrast white, rather than black, discs appearing within the scotoma region. The same dichotomous difference between the two conditions seen in Fig. 3A-C was observed here as well. **(D-F)** Similar to Fig. 3D-F, but with the white stimuli. During expected saccadic inhibition epochs (<100 ms from stimulus onset), there were no differences between the V1 inactivation data (and the presence of a visual stimulus onset in the cortical scotoma) and the no-stimulus data. For monkey A, there was a later reduction in rate in right V1 and a later elevation in rate in left V1, but these effects were always much later than the expected saccadic inhibition time (compare to the intact curves of **B**, **C**). Thus, just like with dark stimuli, saccadic inhibition was rendered statistically unobservable by V1 inactivation. Error bars denote SEM, and all other conventions are like in Fig. 3.

**Figure S6.**
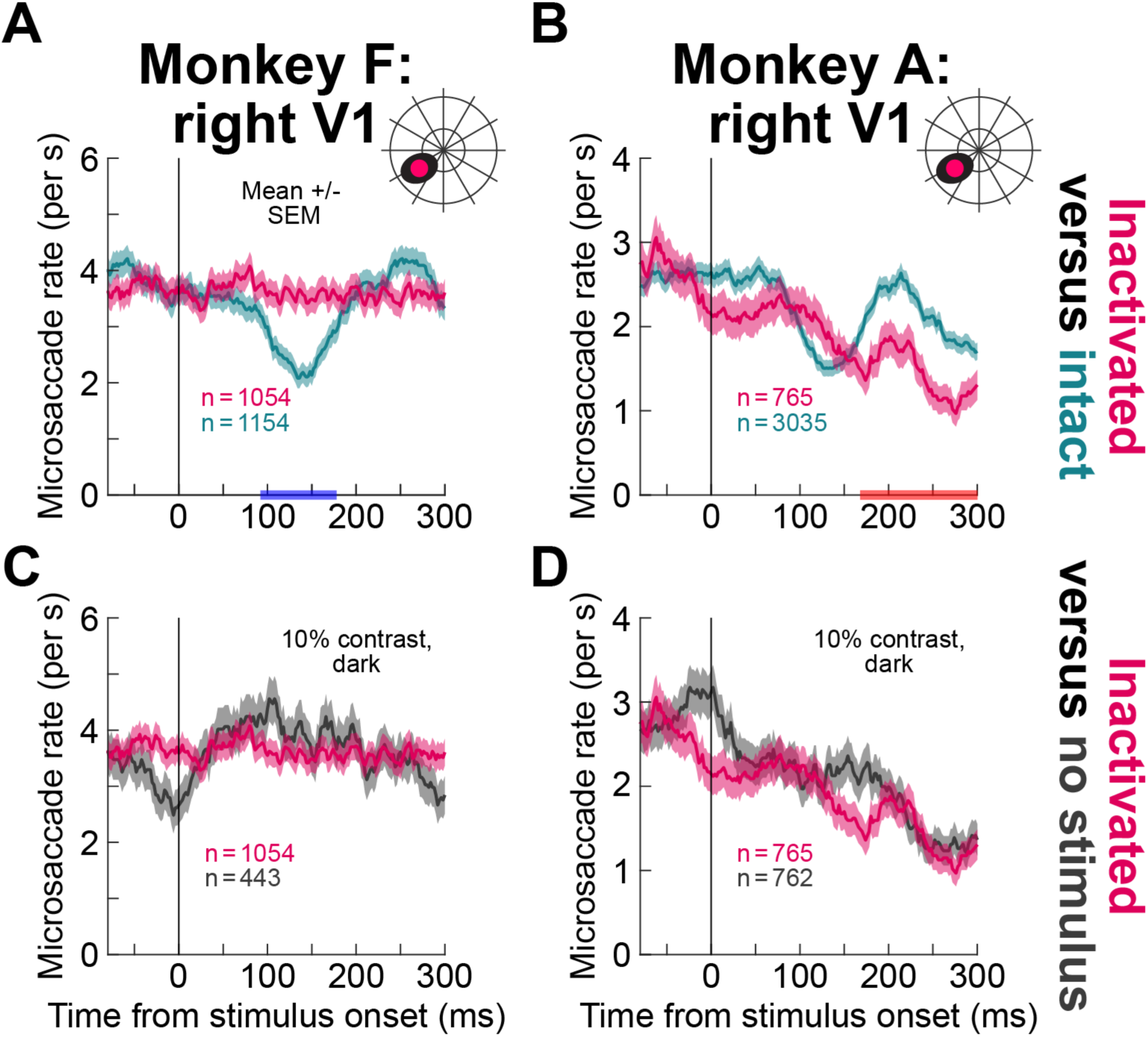
Similar results to. **Fig. 3, but with low contrast visual stimuli. (A, B)** These panels are similar to those in Fig. 3A-C, but now with 10% contrast dark discs. With the stimulus onset being in the scotoma region, the rate curves were not exhibiting saccadic inhibition (reddish curves), whereas they did with an intact V1 (greenish curves). Naturally, the rate modulations in the intact case (greenish curves) were weaker than with 100% contrast stimuli, as expected (81, 89, 117, 118), but they were still present and statistically distinguishable from those with V1 inactivation. **(C, D)** The inactivation curves with the 10% contrast stimuli again did not significantly differ from the no-stimulus curves, also explaining the time-varying inactivation curve of monkey A in **B**. Thus, regardless of stimulus luminance polarity (Fig. S5) or contrast (this figure), the results of Figs. 2, 3 held. Error bars denote SEM, and all other conventions are similar to Fig. 3.

**Figure S7.**
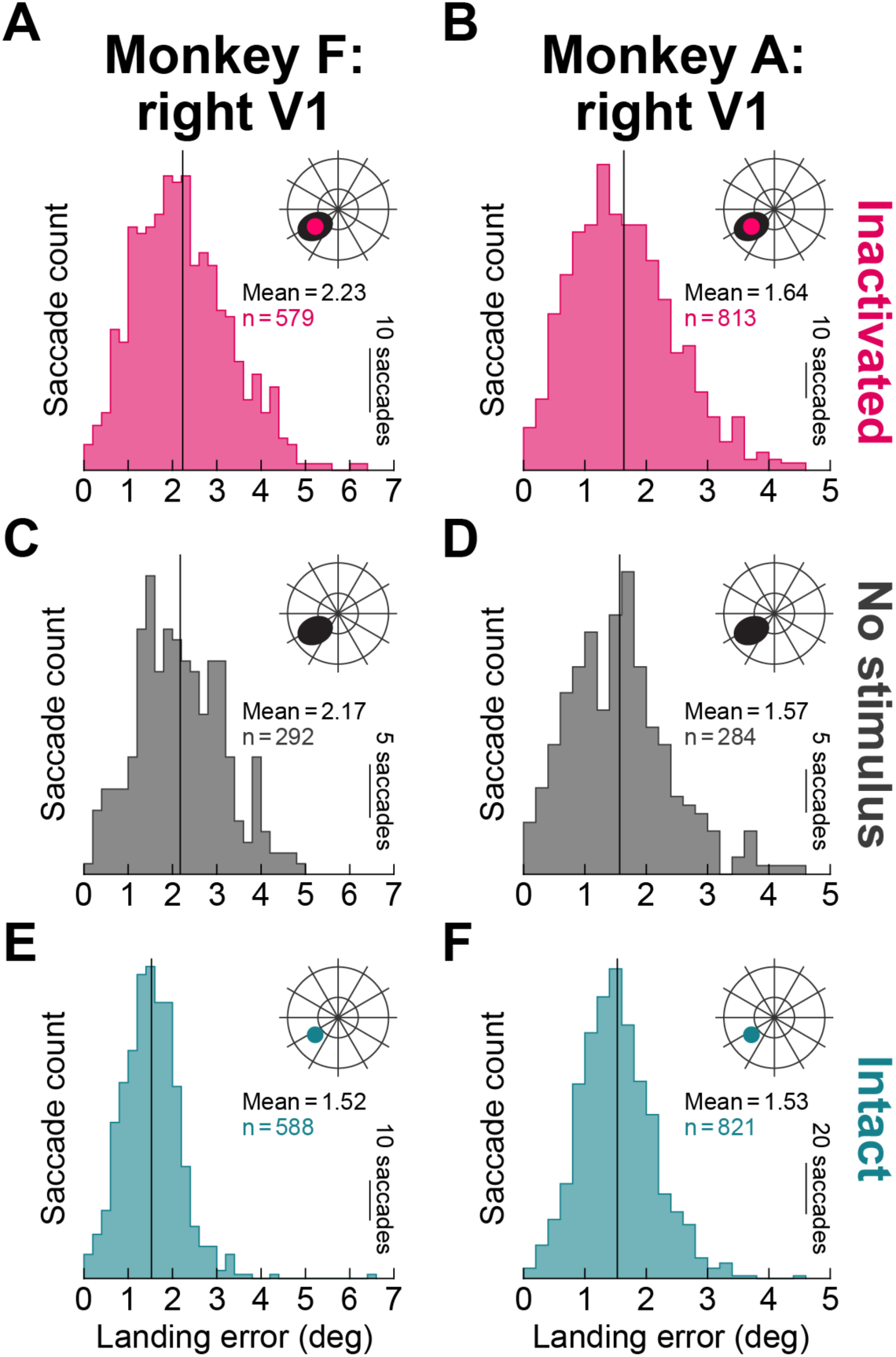
Memory-guided saccade landing errors across V1 inactivation states, and also with no visual onset at all. We trained the monkeys to infer the quadrant of a visual stimulus and generate a memory-guided saccade towards it (Methods). With inactivation (**A**, **B**) or no stimulus (**C**, **D**), the landing errors were of similar order of magnitudes as with an intact V1 and memory guidance for the saccades (**E**, **F**). This suggests that the monkeys were indeed relying on memory in all cases, and paying attention to the affected quadrant during the inactivation periods. Each panel shows a histogram of all saccade landing errors in the memory-guided saccade paradigm.

**Figure S8.**
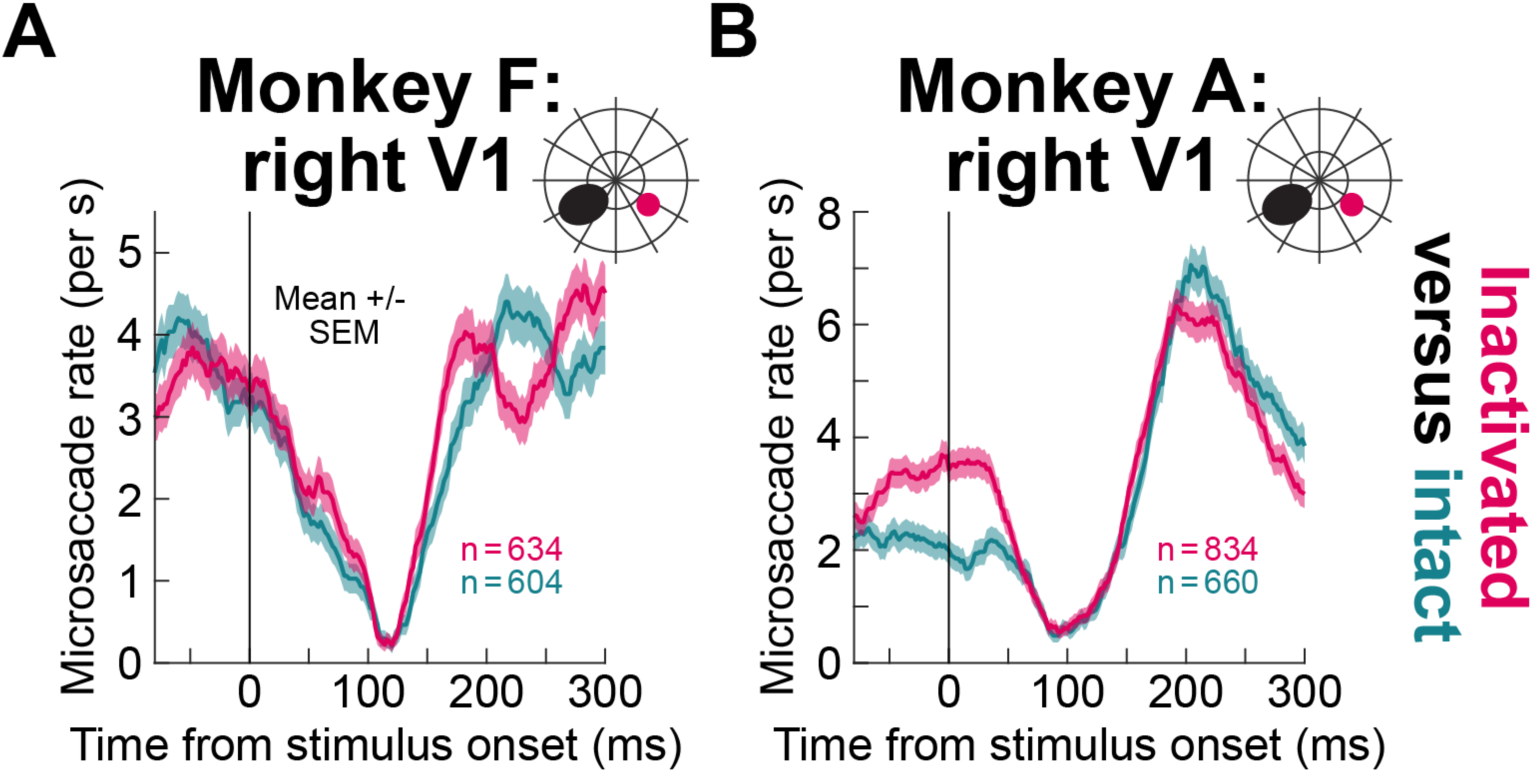
No impairment of saccadic inhibition, in the cognitively demanding task, with stimuli presented in the unaffected visual hemifield. This figure is similar to Fig. S4, but for the memory-guided saccade paradigm. Once again, when the stimulus was presented in the unaffected visual hemifield, saccadic inhibition was not impaired. Monkey A had different pre-stimulus saccade rates in the intact and inactivation conditions, but this did not alter the fact that there was still clear saccadic inhibition in this monkey.

**Figure S9.**
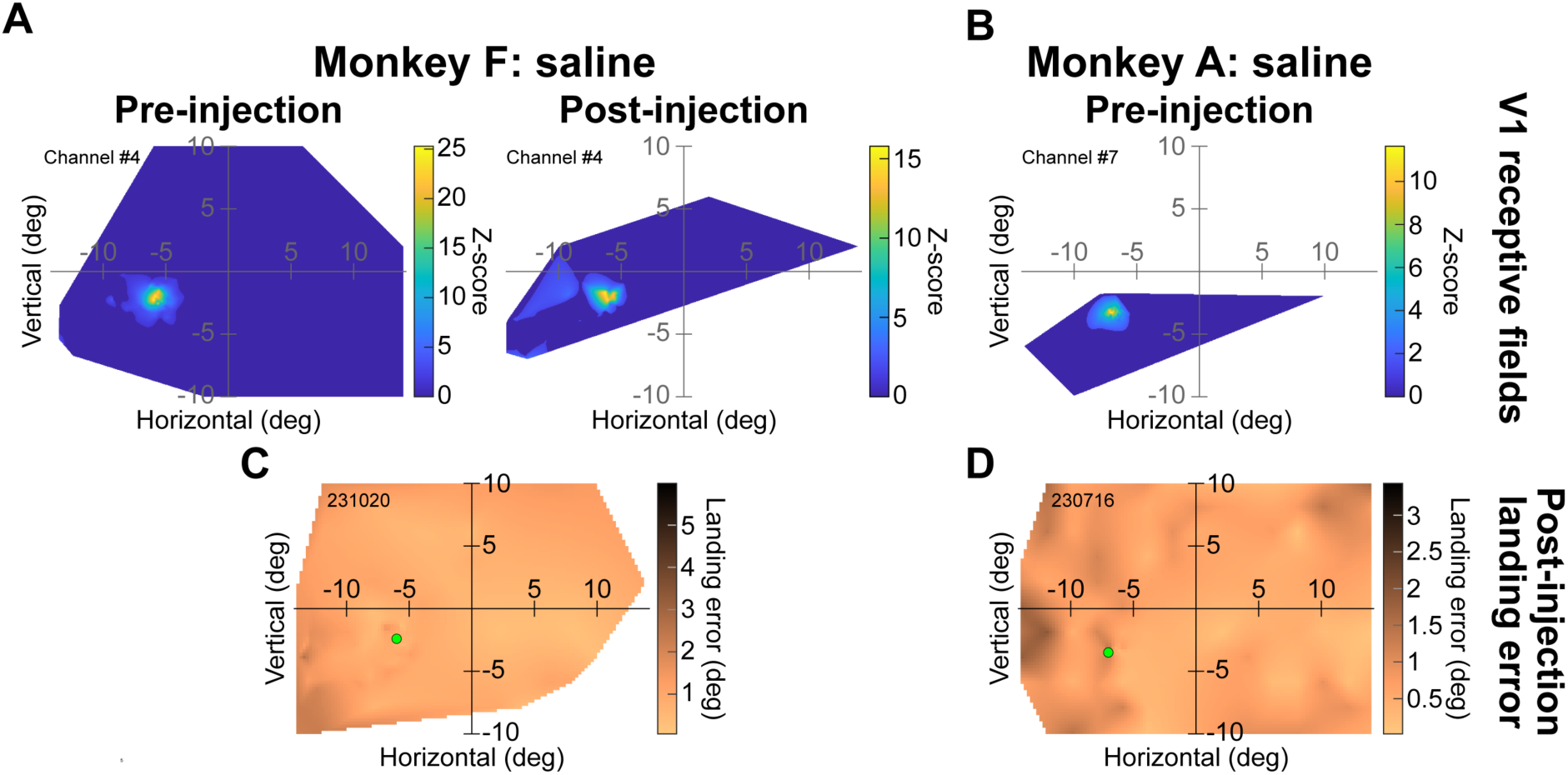
Control saline injections. **(A)** In monkey F, we measured pre-and post-injection visual RF’s like in Fig. 1. We could identify clear, localized V1 RF’s even after saline injection, suggesting that V1 activity was not compromised by saline. **(B)** In monkey A, we retracted the injectrode after the injection period, and thus only measured RF’s in the pre-injection phase. Again, we identified clear, localized V1 RF’s. Thus, for both monkeys, the saline injections were at locations consistent with the muscimol inactivations (Figs. 1, S1). We then checked for potential behavioral deficits in visually-guided eye movements. **(C, D)** Maps of saccade landing errors in the visually-guided saccade task after saline injection. For each session, we used the same z-axis scale as that used for the corresponding muscimol session (for monkey F) or the smallest z-axis scale used with muscimol in Fig. 1 (for monkey A). As can be seen, at the visual stimulus locations targeted by saline injections in each monkey (green discs), there was no elevation in saccadic landing errors relative to other locations. This is very different from the muscimol case (Figs. 1G-I, S1). Thus, the saline injections did not cause a cortical scotoma like muscimol did.

**Figure S10.**
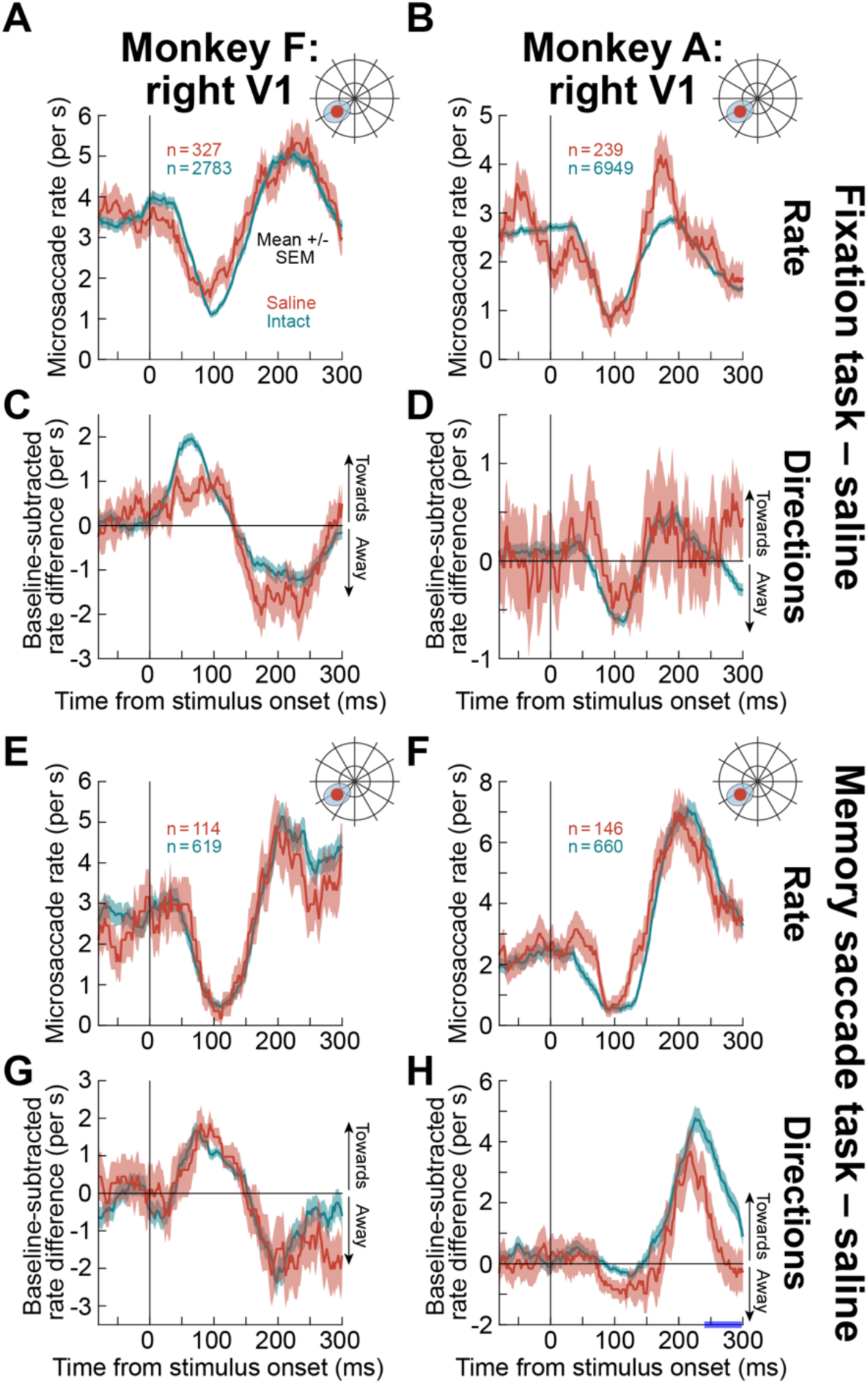
Saccadic inhibition and fixational saccade direction modulations were not impaired with saline injections. (A, **B)** In the same fixation task as that of Figs. 2, 3, we injected saline instead of muscimol in one session for each monkey, and we placed the visual stimulus onset in the corresponding visual field location. Saccadic inhibition still occurred (brownish curves) with similar properties to the intact V1 data (greenish curves). Thus, the muscimol effects on saccadic inhibition were not artifacts of tissue displacement due to fluid injection. **(C, D)** Fixational saccade direction biases (Methods) were also still similar to the intact V1 data. In the latter data, each monkey exhibited a characteristic microsaccade direction oscillation, which replicated earlier behavioral results from the same animals (82) (also see Fig. 8). With the saline injections, the same characteristic oscillations were present in each animal. Thus, both the rate and direction aspects of early stimulus-driven saccadic modulations were not impaired with saline injection. Naturally, the variance in the data was higher in the saline cases because of the lower number of trials. **(E, F)** Same as **A**, **B**, but for the memory-guided saccade task of Fig. 4. Again, saccadic inhibition was intact with the saline injections. **(G, H)** Same as **C**, **D** but for the memory-guided saccade task. The same direction oscillations as with the intact V1 data of each animal were observed. Only at late time intervals in monkey A was there a difference from the intact V1 results (blue epoch on the x-axis in **H**), but these late time intervals are post-inhibition intervals that involve top-down cortical influences (80, 104, 106, 110–112), which could have been different from session to session. Error bars denote SEM.

**Figure S11.**
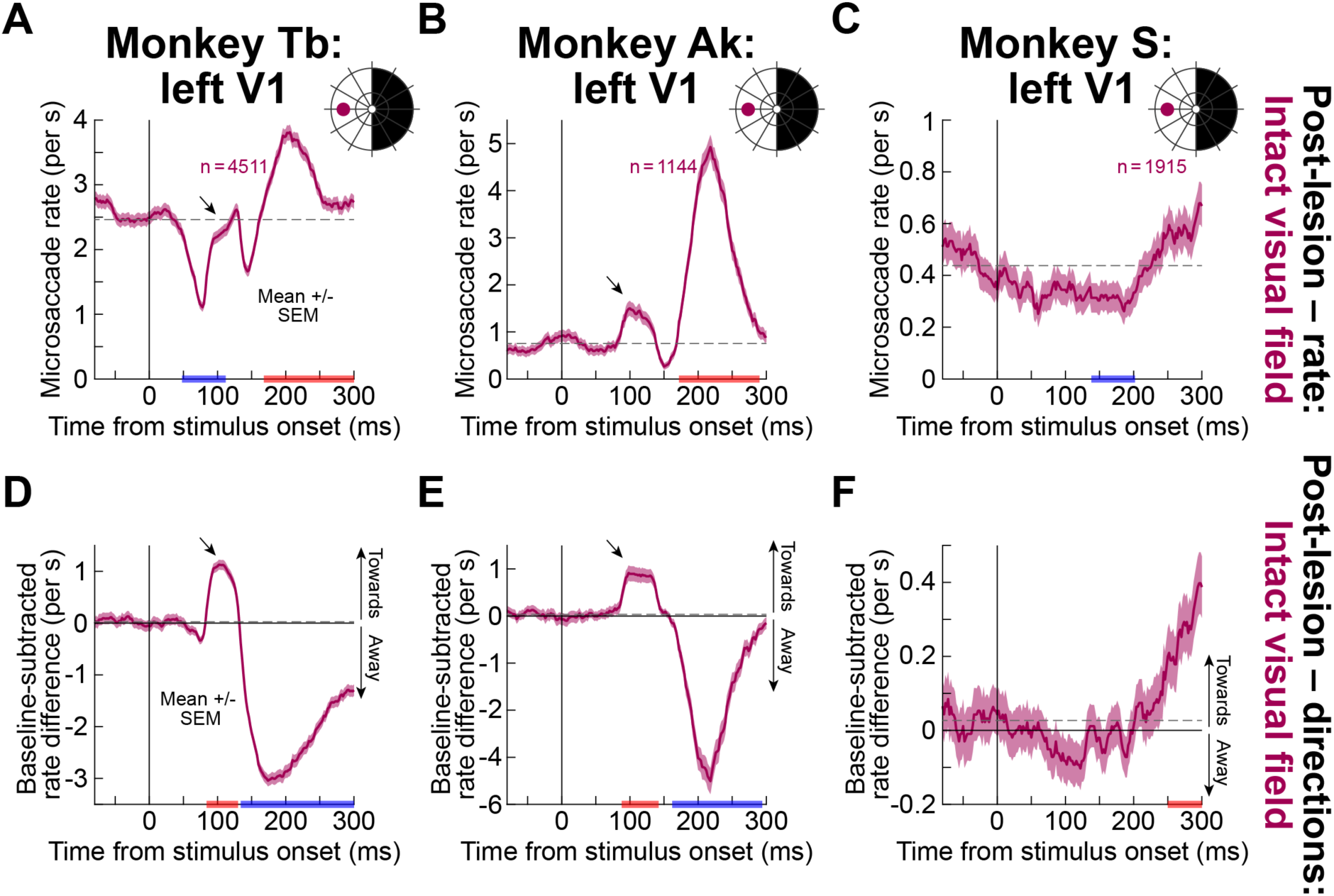
Intact saccadic inhibition for stimuli outside the permanent V1 lesion representation. **(A)** When the visual stimulus appeared in the unaffected visual hemifield after permanent V1 lesions in monkey Tb, there was early saccadic inhibition. Interestingly, there was a rapid burst of fixational saccades right after the onset of the inhibition and before the normally-expected post-inhibition rebound (oblique black arrow). This transient saccade burst was absent for stimuli in the cortical scotoma (Fig. 5) and also absent in pre-lesion measurements (see Fig. S12), and it represents a reflexive orienting of saccade directions towards the stimulus in the intact side (see **D**). **(B)** The second monkey did not show clear early inhibition, but this was likely due to the reflexive saccade burst (oblique arrow), which was also directed towards the appearing stimulus (see **E**) (also see ref. (109) for such reflexes in intact monkeys). Thus, the permanent lesion caused an imbalance that dictates saccade direction oscillations after stimulus onsets (138, 173–177). **(C)** The third monkey exhibited saccadic inhibition for stimuli towards the intact hemifield, similar to monkey Tb. **(D-F)** Direction measures for the fixational saccades in the data of **A**-**C**. For both monkeys Tb and Ak, there were expected saccade direction oscillations, with the early direction bias towards the appearing stimulus being particularly strong (oblique arrows; compare to the pre-lesion measurements in Fig. S12); this suggests an unbalancing of saccade control by a loss of one hemifield representation (177–179). Direction oscillations were less obvious in monkey S (Methods). Error bars denote SEM.

**Figure S12.**
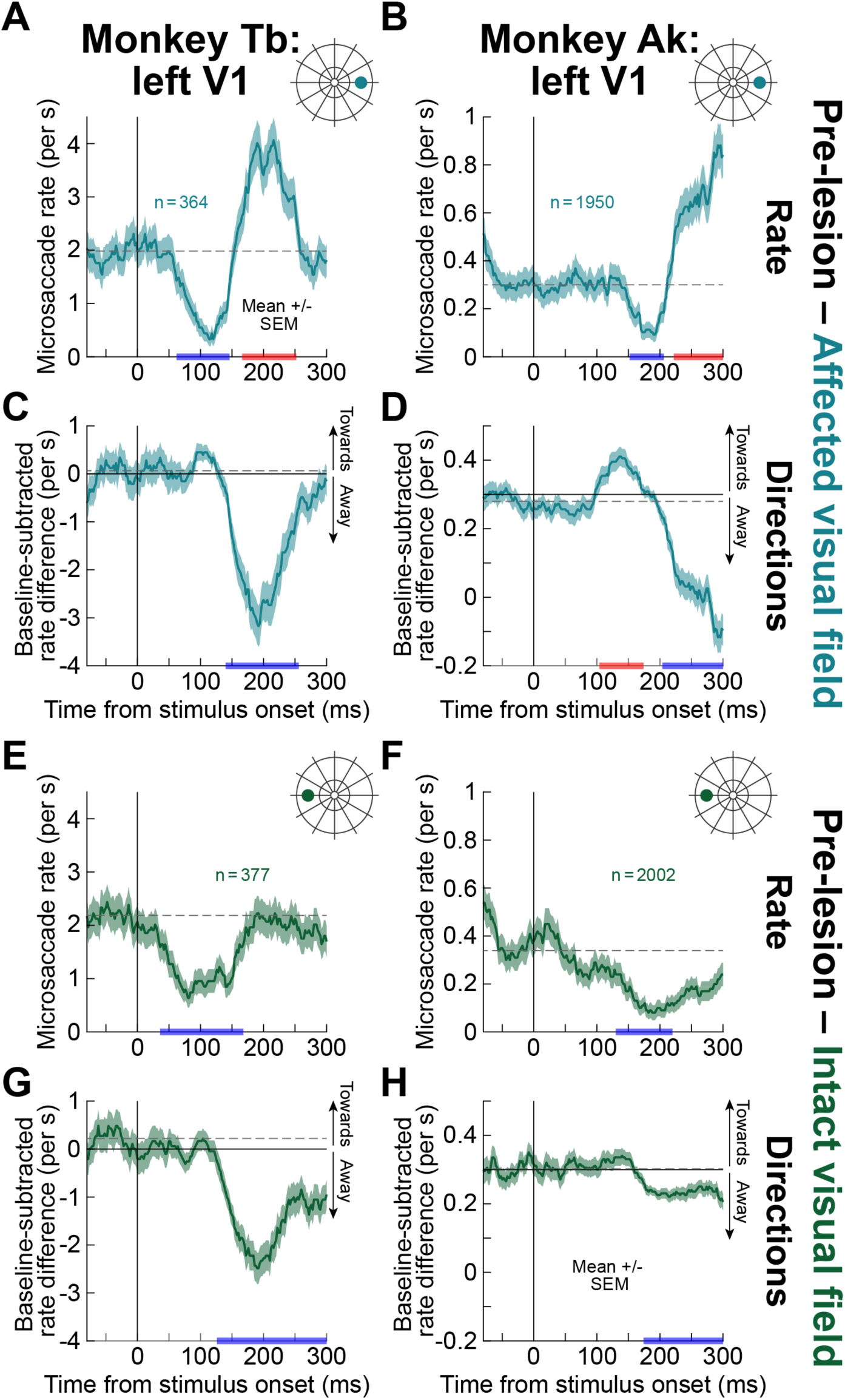
Pre-lesion saccade rate and direction modulations for monkeys Tb and Ak in. **Fig. 5. (A, B)** Both monkeys showed expected saccadic inhibition before the lesion. **(C, D)** Both monkeys also showed expected saccade direction oscillations, although the early bias towards the appearing stimulus did not reach significance for monkey Tb (**C**). **(E-H)** Similar observations for stimuli in the opposite hemifield. The direction biases towards the stimulus location were weaker for this hemifield in both monkeys. Error bars denote SEM.

**Figure S13.**
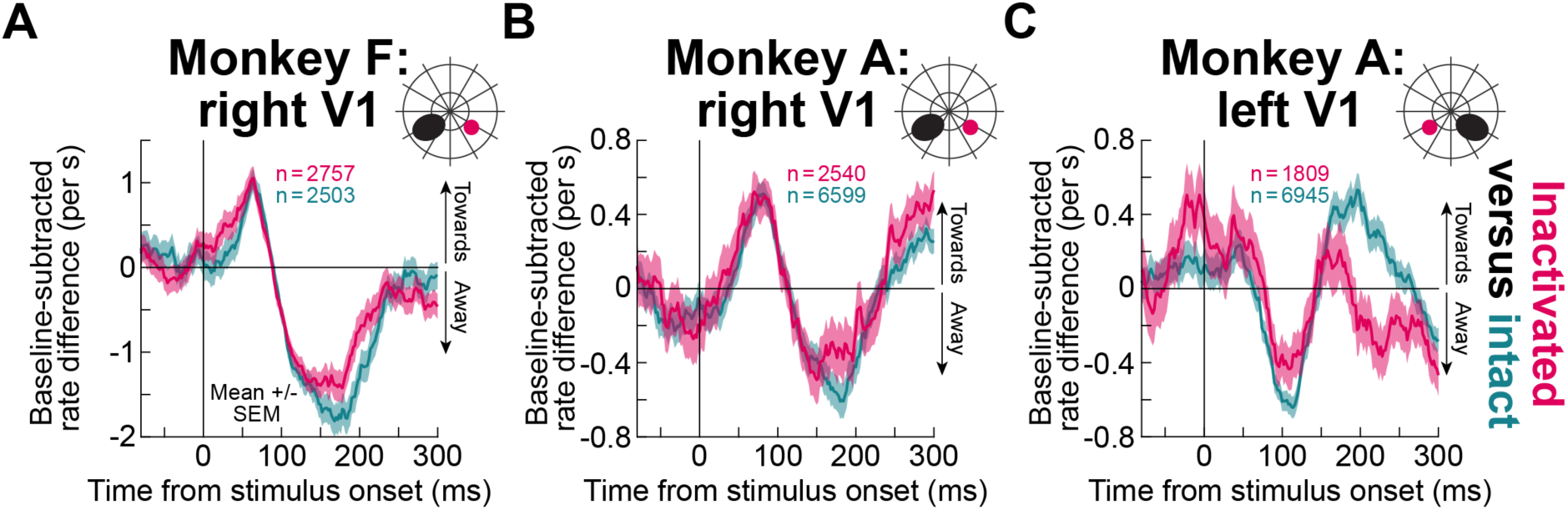
Intact fixational saccade direction oscillations for stimuli outside the affected visual field in the muscimol injection experiments. Like in Fig. 8, we characterized the stimulus-driven fixational saccade direction oscillations of each monkey without (greenish curves; intact) and with (reddish curves; inactivated) reversible V1 inactivation. Here, the stimulus onset was always in the unaffected visual field (inset schematics above each panel). Each monkey’s characteristic direction oscillation was unaltered by V1 inactivation and a stimulus onset in the opposite hemifield. Error bars denote SEM across trials, and all other conventions are like in Fig. 8.

## Supplementary tables

**Table S1.** Results of statistical tests for all figures. In all experimental data, we used cluster-based statistics to identify epochs of statistically significant differences between compared conditions. The table lists all durations of the epochs, as well as the differences in the measured parameters between the conditions. P-values after corrections for multiple comparisons are also included.

| Figure panel | Monkey | Comparison | Time relative to stimulus onset (ms) | Mean difference +/- SEM (microsaccades/s) | P<sub>MC</sub> |
| --- | --- | --- | --- | --- | --- |
| Figure 3 |  |  |  |  |  |
| A | F | Inactivated – Intact | 52-156 | 1.653 +/- 0.0939 | 0 |
|  |  |  | 166-264 | -1.2427 +/- 0.0553 | 0 |
| B | A: right V1 | Inactivated – Intact | 52-130 | 0.9015 +/- 0.0509 | 0.0004 |
|  |  |  | 150-250 | -0.8222 +/- 0.0447 | 0 |
| C | A: left V1 | Inactivated – Intact | 48-128 | 0.8173 +/- 0.0537 | 0.0001 |
|  |  |  | 144-230 | -0.6699 | 0 |
| F | A: left V1 | Inactivated – No stimulus | 232-268 | 0.5248 +/- 0.0142 | 0.0181 |
| Figure 4 |  |  |  |  |  |
| A | F | Inactivated – Intact | 56-168 | 2.0887 +/- 0.0827 | 0 |
|  |  |  | 194-300 | -1.3348 +/- 0.0245 | 0 |
| B | A: right V1 | Inactivated – Intact | 30-146 | 1.8355 +/- 0.0665 | 0 |
|  |  |  | 158-300 | -2.6063 +/- 0.1241 | 0 |
| Figure 5 |  |  |  |  |  |
| A | Tb | Affected – Baseline | 30-206 | -0.7388 +/- 0.0464 | 0 |
|  |  |  | 236-300 | 0.6262 +/- 0.0226 | 0.0008 |
| C | S | Affected – Baseline | 30-184 | -0.2113 +/- 0.0057 | 0 |
| D | Tb | Affected – Baseline | 104-160 | 0.42 +/- 0.0201 | 0.0015 |
|  |  |  | 172-300 | -1.3462 +/- 0.0675 | 0 |
| E | Ak | Affected – Baseline | 96-144 | -0.2558 +/- 0.0061 | 0.0054 |
|  |  |  | 180-234 | -0.2746 +/- 0.0072 | 0.0021 |
| Figure 8 |  |  |  |  |  |
| A | F | Intact – Baseline | 18-124 | 1.1719 +/- 0.0769 | 0 |
|  |  |  | 140-286 | -0.8998 +/- 0.0325 | 0 |
| B | A: right V1 | Intact – Baseline | 64-142 | -0.4756 +/- 0.0297 | 0.0004 |
|  |  |  | 158-208 | 0.3046 +/- 0.0101 | 0.0038 |
| C | A: left V1 | Intact – Baseline | 42-112 | 0.4449 +/- 0.0286 | 0.0003 |
|  |  |  | 130-202 | -0.3173 +/- 0.0135 | 0.0002 |
|  |  |  | 228-300 | 0.3199 +/- 0.0141 | 0.0002 |
| D | F | Inactivated – Intact | 12-122 | -1.1783 +/- 0.0768 | 0 |
|  |  |  | 150-286 | 1.0265 +/- 0.0331 | 0 |
| E | A: right V1 | Inactivated – Intact | 72-126 | 0.4738 +/- 0.0258 | 0.0168 |
|  |  |  | 150-300 | -0.4942 +/- 0.0175 | 0 |
| F | A: left V1 | Inactivated – Intact | 130-220 | 0.5678 +/- 0.0239 | 0.001 |
| G | F | Inactivated – Baseline | 166-216 | 0.4768 +/- 0.0112 | 0.0037 |
| I | A: left V1 | Inactivated – Baseline | 144-300 | 0.3878 +/- 0.0126 | 0 |
| Figure S2 |  |  |  |  |  |
| A | F | Intact – No stimulus | 50-150 | -2.1101 +/- 0.1103 | 0 |
|  |  |  | 198-266 | 1.3138 +/- 0.0494 | 0.0002 |
| B | A: right V1 | Intact – No stimulus | 62-126 | -0.9713 +/- 0.0502 | 0.0009 |
|  |  |  | 162-270 | 0.6883 +/- 0.0192 | 0 |
| C | A: left V1 | Intact – No stimulus | 66-118 | -0.6702 +/- 0.0382 | 0.0026 |
|  |  |  | 158-268 | 0.7326 +/- 0.0232 | 0 |
| Figure S4 |  |  |  |  |  |
| A | F | Inactivated – Intact | 176-220 | 0.5134 +/- 0.0173 | 0.0244 |
| B | A: right V1 | Inactivated – Intact | 92-140 | -0.3693 +/- 0.0216 | 0.0238 |
| C | A: left V1 | Inactivated – Intact | 160-276 | -0.5841 +/- 0.0299 | 0 |
| **Figure S5** |  |  |  |  |  |
| A | F | Inactivated – Intact | 44-152 | 1.3375 +/- 0.0582 | 0 |
|  |  |  | 180-264 | -1.3450 +/- 0.0550 | 0 |
| B | A: right V1 | Inactivated – Intact | 70-180 | 0.9263 +/- 0.0622 | 0.0198 |
|  |  |  | 132-272 | -1.3972 +/- 0.0790 | 0 |
| C | A: left V1 | Inactivated – Intact | 30-112 | 0.9494 +/- 0.0502 | 0.0003 |
|  |  |  | 158-260 | -0.7390 +/- 0.0205 | 0 |
| E | A: right V1 | Inactivated – No stimulus | 132-206 | -0.7961 +/- 0.0261 | 0.0001 |
| F | A: left V1 | Inactivated – No stimulus | 260-290 | 0.5722 +/- 0.0184 | 0.0307 |
| **Figure S6** |  |  |  |  |  |
| A | F | Inactivated – Intact | 92-178 | 1.0467 +/- 0.0434 | 0 |
| B | A: right V1 | Inactivated – Intact | 168-300 | -0.7097 +/- 0.0159 | 0 |
| **Figure S10** |  |  |  |  |  |
| H | A: right V1 | Saline - Intact | 240-298 | -2.2652 +/- 0.0681 | 0.0088 |
| **Figure S11** |  |  |  |  |  |
| A | Tb | Intact – Baseline | 48-112 | -0.6257 +/- 0.0706 | 0.0008 |
|  |  |  | 168-300 | 0.6688 +/- 0.0507 | 0 |
| B | Ak | Intact – Baseline | 172-290 | 2.2076 +/- 0.1649 | 0 |
| C | S | Intact – Baseline | 138-202 | -0.1286 +/- 0.0027 | 0.0019 |
| D | Tb | Intact – Baseline | 84-130 | 0.8498 +/- 0.0547 | 0.0095 |
|  |  |  | 134-300 | -2.1973 +/- 0.0726 | 0 |
| E | Ak | Intact – Baseline | 88-142 | 0.7392 +/- 0.0329 | 0.0019 |
|  |  |  | 162-294 | -1.7134 +/- 0.1325 | 0 |
| F | S | Intact – Baseline | 250-300 | 0.2513 +/- 0.0139 | 0.0036 |
| **Figure S12** |  |  |  |  |  |
| A | Tb | Affected – Baseline | 62-146 | -1.1697 +/- 0.0540 | 0.0002 |
|  |  |  | 166-252 | 1.4354 +/- 0.0655 | 0.0001 |
| B | Ak | Affected – Baseline | 152-206 | -0.1570 +/- 0.0073 | 0.0054 |
|  |  |  | 222-300 | 0.3791 +/- 0.0167 | 0.0002 |
| C | Tb | Affected – Baseline | 140-256 | -2.0690 +/- 0.0990 | 0 |
| D | Ak | Affected – Baseline | 104-174 | 0.1837 +/- 0.0093 | 0.0003 |
|  |  |  | 204-300 | -0.4970 +/- 0.0262 | 0 |
| E | Tb | Intact – Baseline | 36-168 | -1.0914 +/- 0.0353 | 0 |
| F | Ak | Intact – Baseline | 130-220 | -0.2057 +/- 0.0076 | 0.0001 |
| G | Tb | Intact – Baseline | 126-300 | -1.7130 +/- 0.0615 | 0 |
| H | Ak | Intact – Baseline | 174-300 | -0.1356 +/- 0.0026 | 0 |

**Table S2.**
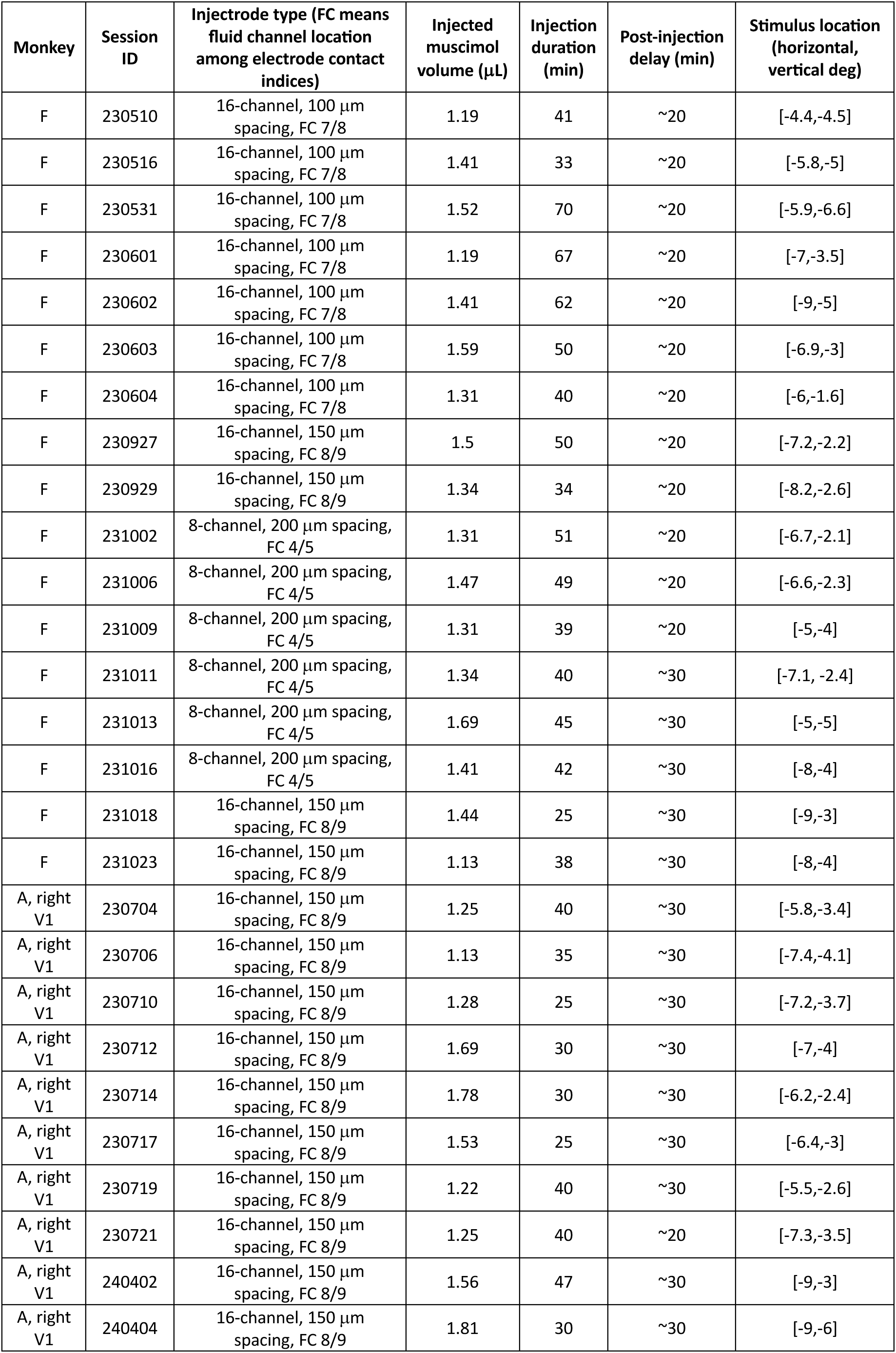
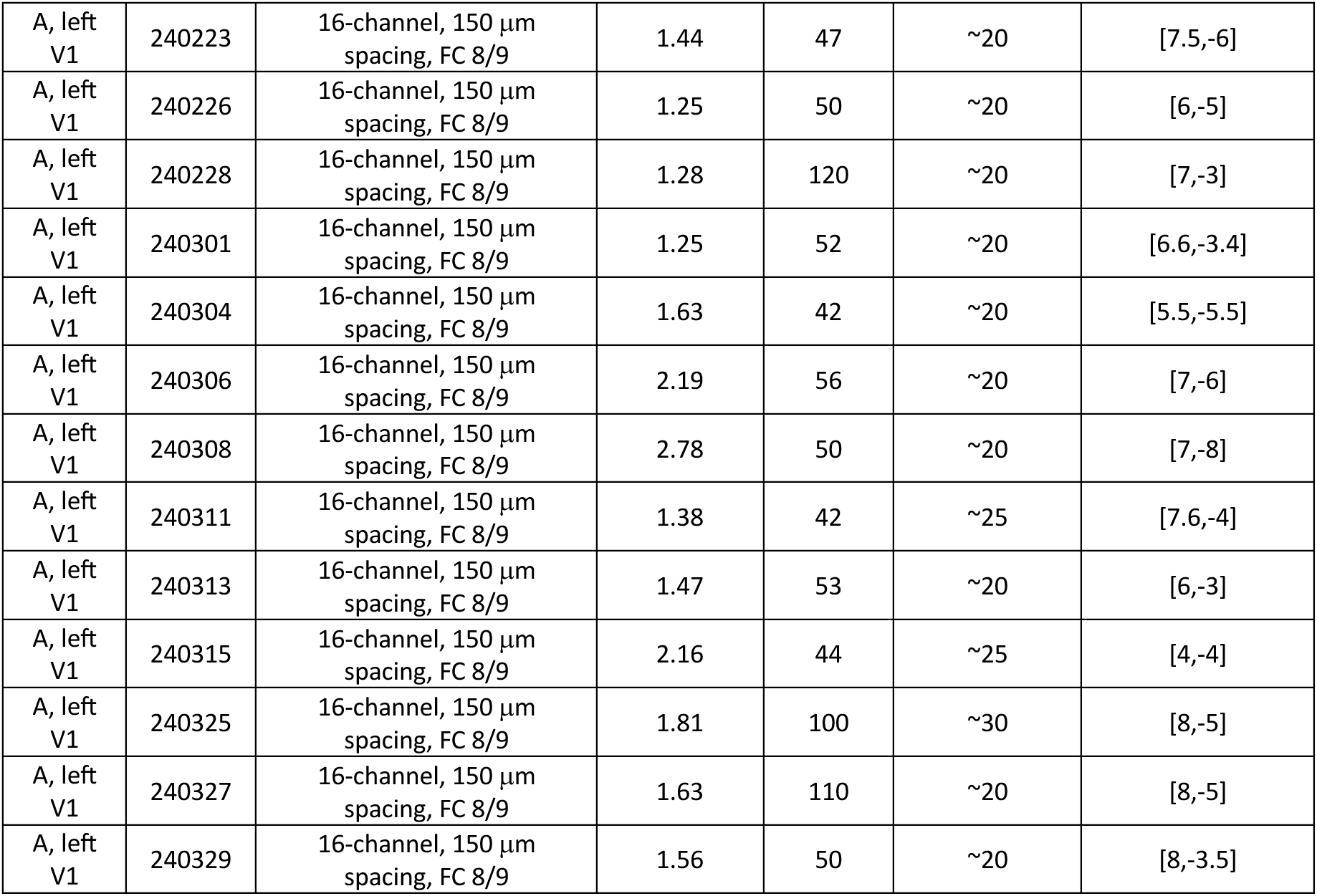
Muscimol injection parameters for all sessions.

| Monkey | Session ID | Injectrode type (FC means fluid channel location among electrode contact indices) | Injected muscimol volume (μL) | Injection duration (min) | Post-injection delay (min) | Stimulus location (horizontal, vertical deg) |
| --- | --- | --- | --- | --- | --- | --- |
| F | 230510 | 16-channel, 100 μm spacing, FC 7/8 | 1.19 | 41 | ~20 | [-4.4,-4.5] |
| F | 230516 | 16-channel, 100 μm spacing, FC 7/8 | 1.41 | 33 | ~20 | [-5.8,-5] |
| F | 230531 | 16-channel, 100 μm spacing, FC 7/8 | 1.52 | 70 | ~20 | [-5.9,-6.6] |
| F | 230601 | 16-channel, 100 μm spacing, FC 7/8 | 1.19 | 67 | ~20 | [-7,-3.5] |
| F | 230602 | 16-channel, 100 μm spacing, FC 7/8 | 1.41 | 62 | ~20 | [-9,-5] |
| F | 230603 | 16-channel, 100 μm spacing, FC 7/8 | 1.59 | 50 | ~20 | [-6.9,-3] |
| F | 230604 | 16-channel, 100 μm spacing, FC 7/8 | 1.31 | 40 | ~20 | [-6,-1.6] |
| F | 230927 | 16-channel, 150 μm spacing, FC 8/9 | 1.5 | 50 | ~20 | [-7.2,-2.2] |
| F | 230929 | 16-channel, 150 μm spacing, FC 8/9 | 1.34 | 34 | ~20 | [-8.2,-2.6] |
| F | 231002 | 8-channel, 200 μm spacing, FC 4/5 | 1.31 | 51 | ~20 | [-6.7,-2.1] |
| F | 231006 | 8-channel, 200 μm spacing, FC 4/5 | 1.47 | 49 | ~20 | [-6.6,-2.3] |
| F | 231009 | 8-channel, 200 μm spacing, FC 4/5 | 1.31 | 39 | ~20 | [-5,-4] |
| F | 231011 | 8-channel, 200 μm spacing, FC 4/5 | 1.34 | 40 | ~30 | [-7.1,-2.4] |
| F | 231013 | 8-channel, 200 μm spacing, FC 4/5 | 1.69 | 45 | ~30 | [-5,-5] |
| F | 231016 | 8-channel, 200 μm spacing, FC 4/5 | 1.41 | 42 | ~30 | [-8,-4] |
| F | 231018 | 16-channel, 150 μm spacing, FC 8/9 | 1.44 | 25 | ~30 | [-9,-3] |
| F | 231023 | 16-channel, 150 μm spacing, FC 8/9 | 1.13 | 38 | ~30 | [-8,-4] |
| A, right V1 | 230704 | 16-channel, 150 μm spacing, FC 8/9 | 1.25 | 40 | ~30 | [-5.8,-3.4] |
| A, right V1 | 230706 | 16-channel, 150 μm spacing, FC 8/9 | 1.13 | 35 | ~30 | [-7.4,-4.1] |
| A, right V1 | 230710 | 16-channel, 150 μm spacing, FC 8/9 | 1.28 | 25 | ~30 | [-7.2,-3.7] |
| A, right V1 | 230712 | 16-channel, 150 μm spacing, FC 8/9 | 1.69 | 30 | ~30 | [-7,-4] |
| A, right V1 | 230714 | 16-channel, 150 μm spacing, FC 8/9 | 1.78 | 30 | ~30 | [-6.2,-2.4] |
| A, right V1 | 230717 | 16-channel, 150 μm spacing, FC 8/9 | 1.53 | 25 | ~30 | [-6.4,-3] |
| A, right V1 | 230719 | 16-channel, 150 μm spacing, FC 8/9 | 1.22 | 40 | ~30 | [-5.5,-2.6] |
| A, right V1 | 230721 | 16-channel, 150 μm spacing, FC 8/9 | 1.25 | 40 | ~20 | [-7.3,-3.5] |
| A, right V1 | 240402 | 16-channel, 150 μm spacing, FC 8/9 | 1.56 | 47 | ~30 | [-9,-3] |
| A, right V1 | 240404 | 16-channel, 150 μm spacing, FC 8/9 | 1.81 | 30 | ~30 | [-9,-6] |
| A, left<br>V1 | 240223 | 16-channel, 150 $\mu\text{m}$<br>spacing, FC 8/9 | 1.44 | 47 | ~20 | [7.5,-6] |
| A, left<br>V1 | 240226 | 16-channel, 150 $\mu\text{m}$<br>spacing, FC 8/9 | 1.25 | 50 | ~20 | [6,-5] |
| A, left<br>V1 | 240228 | 16-channel, 150 $\mu\text{m}$<br>spacing, FC 8/9 | 1.28 | 120 | ~20 | [7,-3] |
| A, left<br>V1 | 240301 | 16-channel, 150 $\mu\text{m}$<br>spacing, FC 8/9 | 1.25 | 52 | ~20 | [6.6,-3.4] |
| A, left<br>V1 | 240304 | 16-channel, 150 $\mu\text{m}$<br>spacing, FC 8/9 | 1.63 | 42 | ~20 | [5.5,-5.5] |
| A, left<br>V1 | 240306 | 16-channel, 150 $\mu\text{m}$<br>spacing, FC 8/9 | 2.19 | 56 | ~20 | [7,-6] |
| A, left<br>V1 | 240308 | 16-channel, 150 $\mu\text{m}$<br>spacing, FC 8/9 | 2.78 | 50 | ~20 | [7,-8] |
| A, left<br>V1 | 240311 | 16-channel, 150 $\mu\text{m}$<br>spacing, FC 8/9 | 1.38 | 42 | ~25 | [7.6,-4] |
| A, left<br>V1 | 240313 | 16-channel, 150 $\mu\text{m}$<br>spacing, FC 8/9 | 1.47 | 53 | ~20 | [6,-3] |
| A, left<br>V1 | 240315 | 16-channel, 150 $\mu\text{m}$<br>spacing, FC 8/9 | 2.16 | 44 | ~25 | [4,-4] |
| A, left<br>V1 | 240325 | 16-channel, 150 $\mu\text{m}$<br>spacing, FC 8/9 | 1.81 | 100 | ~30 | [8,-5] |
| A, left<br>V1 | 240327 | 16-channel, 150 $\mu\text{m}$<br>spacing, FC 8/9 | 1.63 | 110 | ~20 | [8,-5] |
| A, left<br>V1 | 240329 | 16-channel, 150 $\mu\text{m}$<br>spacing, FC 8/9 | 1.56 | 50 | ~20 | [8,-3.5] |

